# The GT1 domain of RNase J ensures RNA quality control through dsRNA binding in Arabidopsis plastids

**DOI:** 10.1101/2025.10.15.682605

**Authors:** Kevin Baudry, Sébastien Skiada, Carmit Burstein, Varda Liveanu, Arnaud Liehrmann, Maddie Ceminsky, Oded Kleifeld, Etienne Delannoy, Alexandra Launay-Avon, Benoît Castandet, Gadi Schuster, David B. Stern

## Abstract

RNase J is a ribonuclease found in bacteria, archaea, and plant chloroplasts, and plays diverse roles in RNA maturation and stability. Chloroplast RNase J is encoded by the nuclear *RNJ* locus and is essential for embryo maturation. Arabidopsis or tobacco plants depleted for RNase J accumulate massive amounts of double-stranded RNA, which interferes with translation and causes chlorosis. Land plant RNase J uniquely contains a C-terminal GT1 domain, a DNA-binding motif found in transcription factors. Here, we have used complementation of an Arabidopsis *rnj* mutant with versions of RNase J with a mutated or deleted GT1 domain to investigate its role in RNase J function. We show that *in vitro*, the recombinant GT1 domain binds both double-stranded RNA and DNA, but not single-stranded nucleic acids, with no sequence specificity. Furthermore, while RNase J lacking GT1 binding complements the *rnj* mutant, these plants accumulate high levels of dsRNA as detected by immunolocalization and RNA-Seq. GT1 mutations also change RNase J solubility *in vivo*, suggesting that the GT1 domain is involved in localization within the plastid. Taken together, our results suggest that the GT1 domain plays a key role in dsRNA removal through localizing the enzyme and/or selectively binding the dsRNA substrate.

**GRAPHICAL ABSTRACT:** 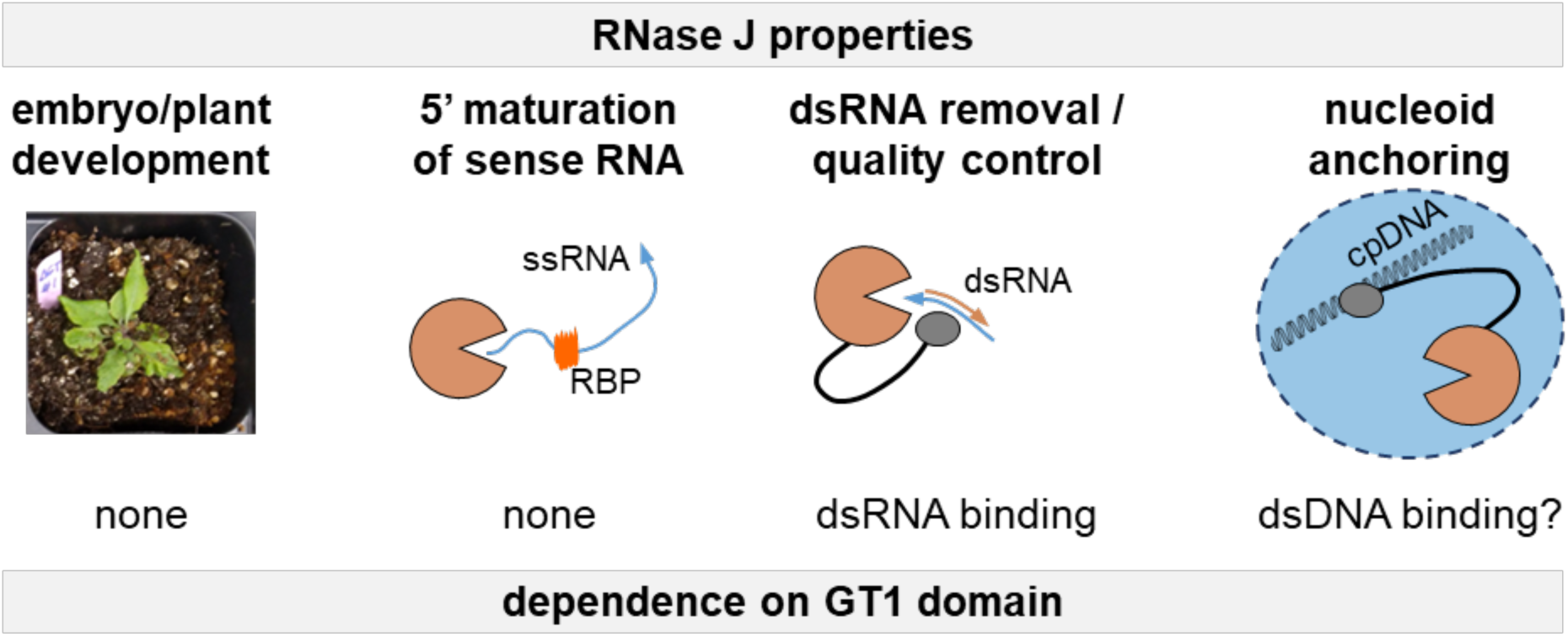

## INTRODUCTION

RNA quality control refers to processes that selectively remove damaged or nonfunctional transcripts that can interfere with normal cellular functions. Efficient RNA quality control ultimately depends on ribonucleases that degrade such transcripts. In plant chloroplasts, RNA quality control is critical because transcription is generally relaxed, with both strands of the ∼150 kb genome fully transcribed from a plethora of promoters. In *Arabidopsis thaliana*, genome-wide studies identified over 200 chloroplast transcription initiation sites (1), consistent with results obtained earlier in barley (2). Systematic analysis of the chloroplast transcriptome, however, shows that a relatively small number of discrete transcripts are retained above background level, mainly corresponding to known genes (3). Therefore, the chloroplast has a robust capacity to cull primary transcripts.

Of the chloroplast ribonucleases studied to date (4), two stand out for the severe effect on plant development when their expression is reduced or eliminated. One is RNase E/G, an endoribonuclease whose absence causes stunted growth and chlorosis, associated with a failure to properly process polycistronic transcripts (5, 6). The other is RNase J (RNJ), which is required for embryo maturation in Arabidopsis (7, 8) and important for seedling development in rice (9), and whose depletion by virus-induced gene silencing in tobacco leads to high accumulation of dsRNA, which interferes with translation of the sense strand (10). Additionally, RNase J plays a role in exonucleolytic 5’ end maturation in cooperation with pentatricopeptide repeat proteins (11), although it is the removal of antisense RNA that is likely its essential function in embryos. Recombinant Arabidopsis RNase J exhibits both endoribonuclease and 5’→3’ exoribonuclease activities (12), which could work cooperatively to efficiently degrade unwanted transcripts and/or complete 5’ end maturation, using the same active site (13).

While RNase J is generally a highly conserved metallo-β-lactamase, orthologues found in land plants stand out for their C-terminal GT1 domain. GT-1 factors are light-responsive transcription factors found in plants which bind to the GT promoter element via a trihelical domain containing three conserved tryptophans (14, 15). The question of why a chloroplast ribonuclease would contain an ostensibly sequence-specific DNA-binding domain prompted the studies described here.

Using *in vitro* binding assays and the generation and analysis of transgenic plants, we show that the GT1 domain is not essential for plant development. Either deletion or mutation of the GT1 domain, however, leads to the accumulation of dsRNA, much of which is derived from noncoding regions of the chloroplast genome, while transcripts from genic regions accumulate to lower levels. The RNJ GT1 domain was found to bind either dsDNA or dsRNA with similar affinity *in vitro*, but without the sequence specificity reported for GT-1 transcription factors. The dsDNA binding activity could be related to localization of RNase J to the plastid nucleoid, which is a diffuse body containing nucleic acids and proteins, as shown in maize and Arabidopsis (16, 17). In contrast, dsRNA binding would be expected to recognize paired sense-asRNAs, where predominantly the asRNA would be targeted for degradation. Our results provide a rationale for recruitment or retention of a GT1 domain in plant RNase J, and further illuminate the mechanism of generalized RNA quality control in the chloroplast.

## MATERIALS AND METHODS

### Genetic material and growth conditions

The T-DNA insertion mutant in At5g63420, *rnj-2* (SAIL_343_B07), was previously described (8). Plants were grown on soil in long day conditions (16h light/8h dark) at 22°C under fluorescent lighting (120-140 µmol.m^-2^.s^-1^).

### Cloning, expression, and purification of recombinant proteins in *E. coli*

cDNA fragments were cloned into pET-28b and designed to create a fusion protein with an N- terminal maltose-binding protein (MBP) and a C-terminal His_6_ tag, inserted between the EcoRI and XhoI restriction sites. Full-length RNJ (RNJ-FL; amino acids 1-911), GT1 alone (GT1; amino acids 817-911) or GT1 containing the upstream CTD-linker (CGT1; amino acids 604- 911) were amplified using the primers listed in Table S1. The ΔCGT1 moiety was created from RNJ-FL by deleting the CGT1 domain (amino acids 604-863) using site-directed mutagenesis. Site-direction mutagenesis of the conserved tryptophans was performed using NEBaseChanger, Q5 Site-Directed Mutagenesis Kit (New England Biolabs), and the corresponding oligonucleotides listed in Table S1.

The resulting expression plasmids were transformed into *E. coli* Rosetta^TM^(DE3) cells. A one-liter culture was grown in LB medium at 37°C with shaking until the OD_600_ reached 0.6, when expression was induced by the addition of 1 mM IPTG, then the culture was incubated for an additional 3 h at 28°C. For protein purification, the bacterial cells were harvested by centrifugation, resuspended in 20 ml of ice-cold lysis buffer (30 mM Tris-HCl, pH 7.7, 150 mM NaCl), and lysed by passage through a microfluidizer operating at 80 psi. The resulting lysate was clarified by centrifugation at 17,000 rpm for 20 min. The soluble lysate was incubated with Ni-NTA agarose beads overnight to bind the His6-tagged protein, washed with at least 10 column volumes of Ni-binding buffer (20 mM sodium phosphate, 0.5 M NaCl, 20 mM imidazole) and then loaded onto a column. The protein was eluted using Ni-binding buffer containing 500 mM imidazole.

Fractions confirmed to contain the CGT1 protein by SDS-PAGE were pooled and diluted 1:1 with amylose buffer (30 mM Tris-HCl, pH 8.0, 300 mM NaCl, 1 mM DTT), and incubated overnight with amylose resin. After washing with at least 10 column volumes of amylose buffer, the bound CGT1 protein was eluted with amylose buffer supplemented with 1% maltose and 5% glycerol. The presence and purity of the final protein product were verified by SDS-PAGE. Due to their inherent instability, purified recombinant proteins were stored in small aliquots at - 80°C in amylose buffer containing 50% glycerol and were utilized in experiments within two weeks of purification.

### Electrophoretic Mobility Shift Assay (EMSA)

dsDNA and dsRNA probes were prepared by annealing 100 µM of complementary oligonucleotides in 80 µl of STE buffer (100 mM NaCl, 10 mM Tris-HCl pH 8.0, 1 mM EDTA). The mixture was heated to 100°C for 5 min and allowed to cool slowly overnight. All probes used are listed in Table S2. Binding reactions were performed in a 10 µl volume containing EMSA reaction buffer (25 mM HEPES pH 7.0, 5% glycerol, 50 mM KCl, 1.1 mM EDTA, 5 mM DTT), 100 nM of the DNA or RNA probe, and the purified protein with incubation for 15 min at 37°C. Before loading the samples, a 6% native polyacrylamide gel (containing 0.5x TBE buffer and 2.5% glycerol) was pre-run in 0.5x TBE running buffer (45 mM Tris-Borate, 1 mM EDTA) at 80 V for 30 minutes. Following the run, the gel was stained with SYBR™ Gold in TE buffer to visualize the nucleic acids.

### ORF cloning, plant transformation, and selection

Sequences encoding RNJ versions flanked by Gateway *att*B recombination sites were synthesized by Twist BioScience (San Francisco, US). WTc corresponds to the full-length Arabidopsis RNJ CDS with no intron and the 2.0 kb region upstream of the ATG start codon, which contains the native promoter and 5’ UTR. mGT corresponds to the WTc construct but with the three GT1 conserved Trp residues (W821A, W850A, W873A) mutated to Ala. ΔGT corresponds to the WTc construct but the GT1 domain is truncated as previously described (12), where amino acids 805 to 904 were removed. ORFs were inserted into pDONR221 using Gateway BP clonase (Invitrogen), then subcloned using LR clonase into pGWB510 (18), which that allows expression from the inserted *RNJ* promoter and adds a C-terminal FLAG tag. Heterozygous *rnj-2/+* plants were transformed using C58C1 pMP90 *Agrobacterium tumefaciens* by floral dip (19) and selected on half MS media containing 33 µg/mL hygromycin B (PhytoTech Labs). Primers used for genotyping are listed in Table S1.

### Pigment content, chloroplast isolation, stroma/thylakoid separation and solubility fractionation

Pigment quantification was performed by 100% acetone extraction on 30 mg fresh rosette leaves of 1-month-old plants as previously described (20). Chloroplasts were isolated from rosette leaves of 4-week-old plants in a 40%/85% PF Percoll step gradient as previously described (16). Purified chloroplasts were snap-frozen for storage and prior to any further use. Stromal and thylakoid fractions were prepared from thawed isolated chloroplasts. After 10 min on ice, the chloroplast lysate was centrifuged for 20 min at 16,000 x g and 4°C. The supernatant was considered to be the stromal fraction, and the pellet the thylakoid fraction.

To test solubility, about 100 mg of ground rosette leaf was suspended in 400 µL in aqueous protein extraction buffer (100 mM Tris pH 7.5, 5 mM EDTA, 2 mM EGTA, 10% glycerol, 1 mM DTT) and incubated for 10 min at 4°C on a wheel. The sample was centrifuged for 10 min at 16,000 x g and 4°C, with supernatant comprising the soluble fraction. The pellet was resuspended in 200 µL of 2X Laemmli buffer supplemented with 15 mg/mL DTT and vortexed for 15 sec, comprising the insoluble fraction.

### Immunoblot analysis

Ground rosette leaf tissue was resuspended in a ratio of 450 µL of 2X Laemmli supplemented with DTT at 15 mg/mL per 100 mg sample. Total proteins were separated in a 10% SDS-PAGE, then transferred onto a PVDF membrane. The FLAG tag was immunodetected with anti-FLAG F3165 (Sigma) at 1:1000 in 5%Milk/TBS-T, HRP conjugated anti-mouse W402B (Promega) at 1:10000 in 5%Milk/TBS-T, and chemiluminescence with Radiance Q (Azure Biosystems).

### Protoplast isolation and immunolocalization

Prior to protoplast isolation, the enzymatic digestion solution (800 mM mannitol, 2 M KCl, 200 mM MES pH 5.7, 1,5% cellulase R10, 0.4% macerozyme R10) was incubated for 10 min at 55°C and then cooled on ice to room temperature. The solution was then supplemented with 0.1% BSA and 1 M CaCl_2_ and filter sterilized (0.45 µm). Protoplasts were isolated from rosette leaves of 4-week-old plants. Fifteen leaves were cut into thin transverse slices and incubated in 10 mL of enzymatic digestion solution in Petri dishes placed in a desiccator in the dark. Vacuum was applied for 30 min, then the leaf material was incubated for 2.5 hr at room temperature in the dark, and the protoplasts finally released with gentle agitation. 10 mL of cold washing buffer (154 mM NaCl, 125 mM CaCl_2_, 5 mM KCl, 2 mM MES pH 5.7) was added and the suspension transferred to fresh tubes on ice through a mesh (35-75 µm). Protoplasts were pelleted by centrifugation (100 g, 2 min, no brake) and resuspended in 5 mL of cold washing buffer. They were then diluted in washing buffer to reach a concentration of 2x10^5^ protoplasts per mL.

Immunolocalization was performed using poly-L-lysine-coated slides (Avantor). Two hydrophobic squares were drawn on each slide with a ReadyProbe Hydrophobic Barrier Pap Pen (Invitrogen). 30 µL of protoplast suspension was applied to each square and allowed to adhere for 30 min at 37°C. The protoplasts were then fixed with absolute ethanol for 20 min and subsequently blocked for 45 min in 1X PBS containing 0.2% Tween20. The fixed protoplasts were incubated overnight at 4°C with the anti-dsRNA monoclonal antibody J2 (Jena Bioscience) diluted 1:500 in 1X PBS supplemented with 1% BSA. Unbound antibodies were removed by washing the slides three times for 10 min in 1X PBS containing 0.2% Tween20. The slides were then incubated for 45 min at 4°C with the secondary antibody AlexaFluor 488 (Invitrogen) diluted 1:2000 in 1X PBS containing 1% BSA. Slides were washed three times for 10 min in 1X PBS containing 0.2% Tween20. Finally, a drop of Mounting Medium with DAPI – Aqueous (Fluroshield, Abcam) was added to cover the protoplasts. Confocal microscopy was carried out using a Zeiss LSM880 operated with ZEN software.

### Protein immunoprecipitation

60 mg of ground rosette leaf were resuspended in 1 mL of cold Protein Extraction Buffer (1M Tris-HCl pH 7.5, 50% Glycerol, 500 mM EDTA, 100 mM EGTA, 1 tablet of protease Inhibitor EDTA-free [ThermoScientific]) per 100 mL buffer, and incubated for 30 min at 4°C with rotation. Anti-FLAG antibody F1804 (Sigma) was added to a final dilution of 1:2000 and the mixture was incubated overnight at 4°C with rotation. Cellular debris was cleared by centrifugation (30 sec at 8200xg at 4°C) and the supernatant directly loaded on 25 µL of Dynabeads Protein G (Invitrogen) previously washed 2x5 min in 500 µL TBS) and incubated for 1h at 4°C with rotation. Beads were washed 3x5 min with 500 µL cold TBS and resuspended in 500 µL NucleoZOL (Macherey-Nagel) supplemented with 175 µL of nuclease-free water, and RNA was extracted according to the manufacturer’s instructions.

### dsRNA immunoprecipitation

50 mg of ground rosette leaf were resuspended in 750 µL of cold dsRIP buffer (50 mM Tris-HCl pH 7.4, 100 mM NaCl, 3 mM MgCl_2_, 0.5% IGEPAL CA-630, ½ tablet of protease Inhibitor EDTA-free [ThermoScientific] per 15 mL buffer, and 100 u/mL RNasin Ribonuclease Inhibitor [NEB]). The homogenate was vigorously mixed and incubated for 30 min on ice before centrifugation at 20,000 x g for 10 min at 4°C. For the IP, 500 µL of the supernatant was incubated with 5 µL undiluted antibody J2 overnight at 4°C with gentle rotation. The IP solution was then added to 25 µL of Dynabeads Protein G (Invitrogen) previously washed 2x2 min in 500 µL cold dsRIP buffer and incubated for 1h at 4°C. Beads were washed 4x 2 min with 500 µL cold dsRIP buffer and resuspended in 500 µL of NucleoZOL supplemented with 175 µL of nuclease-free water. RNAs from the IP and input fractions were extracted according to the manufacturer’s instructions and quality checked with an Agilent RNA Nano Chip for input and Agilent RNA Pico Chip for IP on an Agilent 2100 Bioanalyzer.

### Total RNA extraction, cDNA synthesis, 5’ RACE, RT-qPCR and dot blots

Leaf total RNA was extracted using the NucleoZOL protocol (Macherey-Nagel) and RNA quality was checked by analyzing 4 µg in a 1.2% formaldehyde-agarose gel. For 5’ RACE, an RNA adaptor was ligated to 2 µg RNA using T4 RNA ligase (NEB) as described (11). SuperScript III (Invitrogen) was used to synthesize cDNA according to the manufacturer’s protocol. 5’ termini were amplified with Gotaq (Promega), using an adapter-specific primer and a gene-specific primer. RT-PCR products were separated in 2% agarose gels. Gene expression was measured by RT-qPCR using iTaq Universal SYBR Green Supermix (BioRad). For each plant, the mean expression level of three technical RT-qPCR replicates was normalized with the mean of the reference gene *Actin1*. All primers used are listed in Table S1. For dot blots, 1 µg of RNA was spotted onto a Hybond-N+ membrane (Cytiva) optimized for nucleic acid transfer. The RNA was crosslinked in a Stratalinker 2400 UV Crosslinker on both sides of the membrane (1.200 µJ x100; 25-50 sec). The membrane was washed in dsRIP lysis buffer (100 mM NaCl; 50 mM Tris-HCl pH 7.4, 3 mM MgCl_2_ 0,5% IGEPAL CA-630) for 5 min with gentle shaking and then incubated in blocking buffer (1X PBS, 0.02% Tween20, 5% nonfat milk) for 1h with gentle shaking. The membrane was then incubated overnight at 4°C with antibody J2 (Jena Bioscience) diluted 1:3,000 in PBS containing 0.02% Tween20 and 5% nonfat milk, with gentle shaking. The membrane was washed 3x5 min in dsRIP lysis buffer and incubated for 2h with goat anti-mouse IgG conjugated with HRP (Sigma-Aldrich) diluted 1:10,000 in PBS/0.02% Tween20/5% nonfat milk), followed by 4x 5 min washes in dsRIP lysis buffer. Detection was performed using the AgriseraECL Super Bright protocol (Agrisera). Chemiluminescence signals were acquired using a ChemiDoc MP imaging system (Bio-Rad) operated with Image Lab software.

### Library preparation, RNA-Seq and Data Analysis

Sequencing libraries were constructed using the POPS platform (IPS2) from 300 ng of total RNA or 15-60 ng of immunoprecipitated RNA (see dsRNA immunoprecipitation section) using the Illumina TruSeq Stranded Total RNA with Ribo-Zero Plant kit (Illumina) with half reaction volumes. Library quality was checked using the Agilent High Sensitivity DNA Chip on the Agilent 2100 Bioanalyzer. Final quantification was performed using the Quant-iT PicoGreen dsDNA kit (Invitrogen) to generate an equimolar pool. Libraries were sequenced (2x150) on a NovaSeq X system (Illumina) by Biomarker Technologies GmbH, Münster, to obtain approximately 30 and 10 million reads per total or IPed samples, respectively. Adapter sequences and bases with a quality lower than a Q-score of 20 were removed using Trimmomatic software (v.36) (21), only keeping reads longer than 30 bases after trimming.

RNA-Seq data mapping, counting, splicing, editing and DEG analysis were conducted with pipelineOGE (default parameters) (22) using four biological replicates. For splicing extent analysis, introns with at least 6 spliced reads found across samples were analyzed. For editing extent analysis, positions with at least 6 edited reads found across samples and an average extent of ≥0.5% were analyzed. For both editing and splicing analyses, p-values were adjusted by the Bonferroni correction method, and considered significant when the q-value was <0.05. For DEG identification in each pairwise comparison, genes with at least 1 CPM (count per million) were kept for TMM normalization and analysis; p-values were adjusted by the Benjamini-Hochberg (BH) procedure and considered significant when the q-value was <0.05. Enrichment analysis was conducted using DiCoExpress (23, 24). Enrichment was performed to identified overrepresented terms among the entire DEG list, or upregulated (log2FC>0, q-value<0.05) or downregulated (log2FC<0, q-value<0.05), and considered significant when the BH-adjusted p-value was <0.05. Three biological replicates of dsRIP-Seq data were obtained and mapped with pipelineOGE. All coverage analysis by segmentation was conducted using the DiffSegR package (25). The significance threshold was set at 0.01 for total RNA analysis, and 0.05 for dsRIP-Seq analysis. To identify *trans*-dsRNA, DiffSegR was used to segment the log ratio of the plus over minus strand. Regions lacking differential strand coverage (p-value >0.1) and with an average length normalized coverage exceeding 10 were considered to be *trans*-dsRNAs.

### Proteomics, protein extraction and digestion, LC-MS/MS and MS data analysis

Two-week-old seedlings were lysed in conditioned medium (8.5 M urea, 400 mM ammonium bicarbonate, and 10 mM DTT), sonicated twice (90%, 10-10, 5 min), and centrifuged (10,000 × *g*, 10 min). The protein amount was estimated using Bradford readings. The samples from the conditioned media were reduced at 60 °C for 30 min, modified with 35.2 mM iodoacetamide in 100 mM ammonium bicarbonate (room temperature for 30 min in the dark), and digested in 1.5 M urea, 66 mM ammonium bicarbonate with modified trypsin (Promega) overnight at 37 °C in a 1:50 (M/M) enzyme-to-substrate ratio. An additional digestion with trypsin was performed for 4 h at 37°C in a 1:100 (M/M) enzyme-to-substrate ratio. The tryptic peptides were desalted using a homemade C_18_ stage tip, dried and resuspended in 0.1% formic acid.

2 µg tryptic peptides were analyzed by LC-MS/MS on an Exploris 480 Orbitrap mass spectrometer (ThermoFisher Scientific) interfaced with a Vanquish Neo UHPLC (ThermoFisher Scientific). Peptides were loaded in solvent A (0.1% formic acid in LC-MS grade water) onto a homemade capillary column (30 cm length, 75 µm inner diameter) packed with Reprosil C_18_- Aqua. Peptides were separated using a linear gradient from 6 to 34% solvent B (80% acetonitrile, 0.1% formic acid in LC-MS grade water) over 180 min, followed by an increase to 100% solvent B in 1 min, and re-equilibration in 100% solvent B for 11 min, at a constant flow rate of 0.15 µl/min.

The mass spectrometer was operated in data-dependent acquisition mode with positive ionization. Full MS scans were acquired in the Orbitrap at a resolution of 120,000 (at m/z 200) with an AGC target set to 300% over an m/z range of 350–1200. The 30 most abundant precursor ions with charge states 2 to 6 were selected for fragmentation. MS/MS scans were acquired at a resolution of 15,000 using an isolation window of 1.3 m/z, normalized collision energy of 27% (HCD), The intensity threshold for MS/MS triggering was set to 8 × 10³, and dynamic exclusion was applied for 30 s.

Data analysis was conducted using FragPipe v. 23.0 (https://fragpipe.nesvilab.org/) in DDA+ mode with the default label-free quantification and match-between-runs (LFQ-MBR) workflow. Searches were performed with trypsin specificity allowing up to two missed cleavages. Carbamidomethylation of cysteine was set as a fixed modification, while methionine oxidation and protein N-terminal acetylation were specified as variable modifications. Spectra were searched against the Arabidopsis thaliana UniProt reference proteome (Taxon ID 3702, downloaded Jan 5, 2025; 39,396 sequences) supplemented with common contaminants and decoy sequences. Identifications were filtered to a 1% false discovery rate (FDR) at peptide and protein levels. Subsequent data processing and statistical analysis were conducted in Perseus v2.1.4 (https://maxquant.net/perseus/). Raw MaxLFQ intensities were log₂- transformed, and proteins were retained if they contained MaxLFQ intensity values in at least three replicates within one experimental group. Statistical testing was performed using a two-sample t-test with permutation-based FDR control (FDR = 0.05, s₀ = 0.1) under default settings.

## RESULTS

### Land plant RNJs contain a GT1 domain

RNase J is a conserved endo- and exoribonuclease found in all three domains of life. The metallo-β-lactamase (MBL) and β-CASP domains form the conserved nuclease domain containing several conserved signature motifs (Fig. 1A, Supplemental Fig. S1A). RNase J usually contains a C-terminal domain (CTD) thought to be required for enzyme oligomerization in bacteria, and sharing structural similarity with the Small Domain of RNase E also involved in its oligomerization (26–29). The sequence of this core enzyme (MBL-β-CASP-CTD) is conserved across orthologs apart from archaea, which lack the CTD.

**Figure 1.**
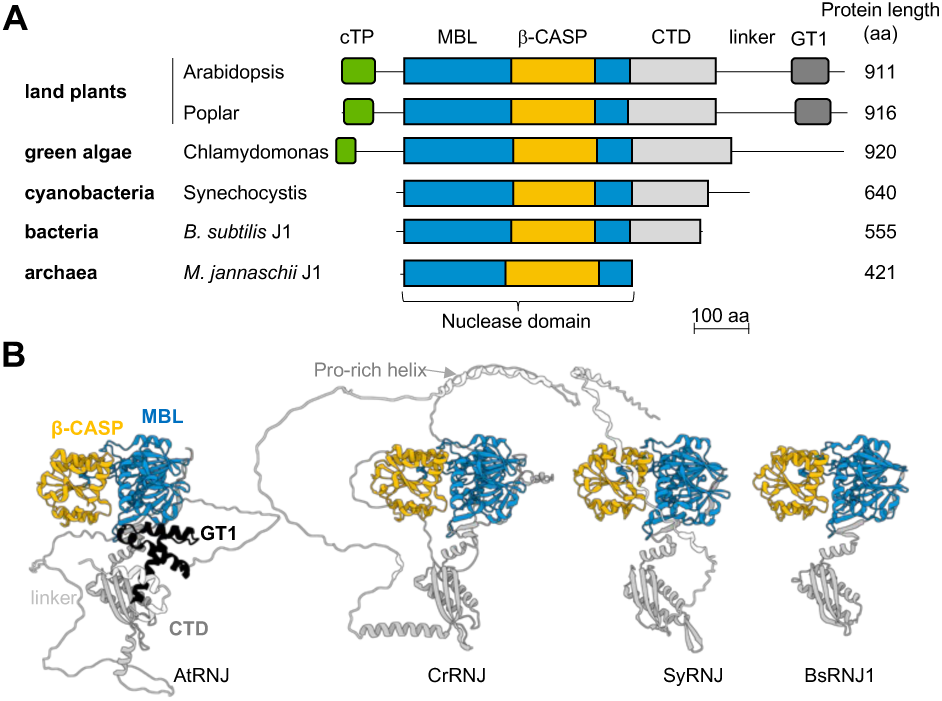
RNase J protein domains and structure. **(A)** RNJ linear structures from *Arabidopsis thaliana*, *Populus trichocarpa*, *Chlamydomonas reinhardtii*, *Synechocystis* sp. PCC 6803, *Bacillus subtilis*, and *Methanocaldococcus jannaschii*. Domains drawn to scale are the cTP, chloroplast target peptide; MBL, Metallo-β-lactamase domain; β-CASP, β-CASP domain; CTD, C-terminal domain; GT1, GT1 domain. On the right, protein length is indicated. **(B)** AlphaFold predicted structures as accessible from the AlphaFold Protein Structure Database (https://alphafold.ebi.ac.uk/). Arabidopsis (At, AF-Q84W56-F1); Chlamydomonas (Cr, AF-B2YFW5- F1); Synechocystis (Sy, AF-P54123-F1); and Bacillus (Bs, AF-Q45493-F1). Domains are colored as in panel A and as labeled for AtRNJ. The N-terminal region upstream of the MBL domain is hidden for easier viewing. The proline-rich helix unique to the *Chlamydomonas* linker region is indicated.

Structural predictions for the core enzyme made by AlphaFold (https://alphafold.ebi.ac.uk/) agree with crystallography-based structures and show an overall conserved structure and shape with a few discrepancies (Supplemental Fig. S1B-C). Some forms of RNase J have an extended C-terminal, such as in the cyanobacterium *Synechocystis* PCC6803, *Chlamydomonas*, and land plants. While some of these C-terminal extensions do not encode known domains or predict conserved folding, in land plants this extension contains a conserved domain, GT1 (Fig. 1B, Supplemental Fig. S1B-C). Initially described in the pea transcription factor GT-1, this domain consists of a trihelical structure and binds to DNA (14, 30). In the following experiments, we investigated the binding properties of the GT1 domain derived from Arabidopsis RNJ.

### The RNJ GT1 domain binds non-specifically to double stranded nucleic acids *in vitro*

A previous investigation showed that the GT1 domain of Arabidopsis chloroplast RNase J is not required for *in vitro* nuclease activity (12). Since this domain is found in transcription factors where it binds sequence-specifically to promoters of some light-responsive genes (14), we investigated the ability of recombinant CGT1, which contains part of the CTD along with the GT1 domain (Supplemental Fig. S2A), to bind DNA. In transcription factors, the GT1 domain recognizes a degenerate purine-rich sequence called the GT element, which for the Arabidopsis GT-1 transcription factor is GGTTAA (31). A double-stranded DNA probe containing four tandem repeats of a 14 bp sequence containing this element, hereafter called 4GT, was used to assess the binding ability of CGT1. Gel shift assays showed robust binding of 4GT with a K_D_ of 298nM (Fig. 2A). In parallel, we tested its ability to bind single-stranded DNA with the 4GT sequence, and none was observed, suggesting specificity for dsDNA (Fig. 2B). The binding abilities of two other proteins were tested, namely full-length Arabidopsis RNJ and ΔCGT1, the latter containing solely the nuclease domain and CTD (Supplemental Fig. S2A). Like CGT1, full-length RNJ was able to bind 4GT dsDNA, however, neither ΔCGT1 nor the maltose binding protein (MBP) tag showed any binding activity (Fig. 2C-D, Supplemental Fig. S2B). These results verify that the CGT1 domain is both necessary and sufficient to impart DNA-binding activity *in vitro*.

**Figure 2.**
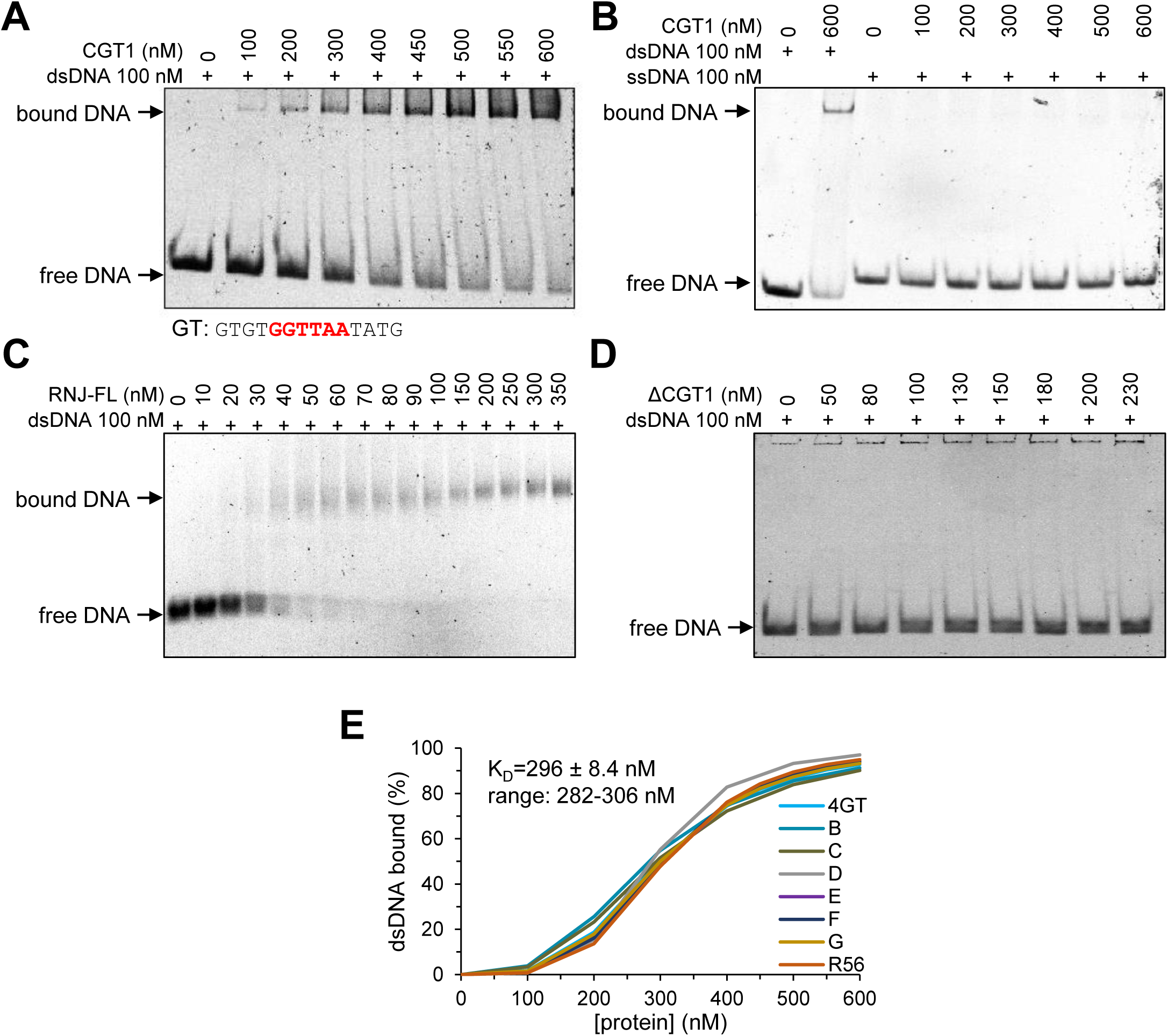
The recombinant GT1 domain binds dsDNA without sequence specificity. **(A)** Gel shift assay using a recombinant maltose binding protein-GT1 fusion and a 56 bp DNA probe with four tandemly repeated consensus GT sequences, as shown below the gel with the invariant GGTTAA motif in red. **(B)** As in panel **A**, except the DNA probe was single-stranded. The dsDNA probe was used as a positive control in the second lane from left. **(C)** As in panel **A**, except recombinant full-length RNJ was used. **(D)** As in panel **A**, except recombinant ΔCGT1 was used. **(E)** Binding curves obtained by quantification of the stained gels assaying the various dsDNA probes (corresponding to panel **A** and Supplemental Fig S2, panels H-N). The sequences for probes B to G are given in Supplemental Table S2. K_D_ values calculated from each curve are shown. The average K_D_ value is given with standard deviation.

To further characterize dsDNA binding ability length variation of the probe was tested, using probes 1GT to 5GT. Although a single element was not bound, the other probes bound with affinities increasing with the length of the probe (Supplemental Fig. S2C-G), consistent with previous results obtained with the GT1 transcription factor (32). Since the transcription factor displayed sequence-specific DNA binding within the GT elements (32), we tested whether this is true for the RNJ GT1 domain. To do so, we used probes that varied within the GT element and/or surrounding nucleotides, as well as a random non-repeat sequence. In contrast to the characterized transcription factor and despite the similarity of the amino acid sequence and the predicted structure, recombinant CGT1 bound all tested probes with strikingly similar affinities (Supplemental Fig. S2H-O). Therefore, CGT1 appears to lack the sequence specificity of its transcription factor counterparts.

While CGT1 was expected to bind DNA, RNJ could potentially use this domain to help target transcripts if it could bind RNA in addition to DNA. To test this possibility, CGT1 was used in gel shift assays with single- and double-stranded RNA (Fig. 3, Supplemental Fig. S3). These experiments showed that CGT1 can bind various randomized dsRNA sequences (Fig. 3A), with an affinity marginally higher than that measured for dsDNA, and with longer probes binding more tightly than shorter ones (Fig. 3B), as was observed with DNA. Single-stranded RNA, however, did not bind CGT1 (Fig. 3C). Therefore, Arabidopsis CGT1 binds double-stranded nucleic acids nonspecifically, but does not bind to single-stranded nucleic acids.

**Figure 3.**
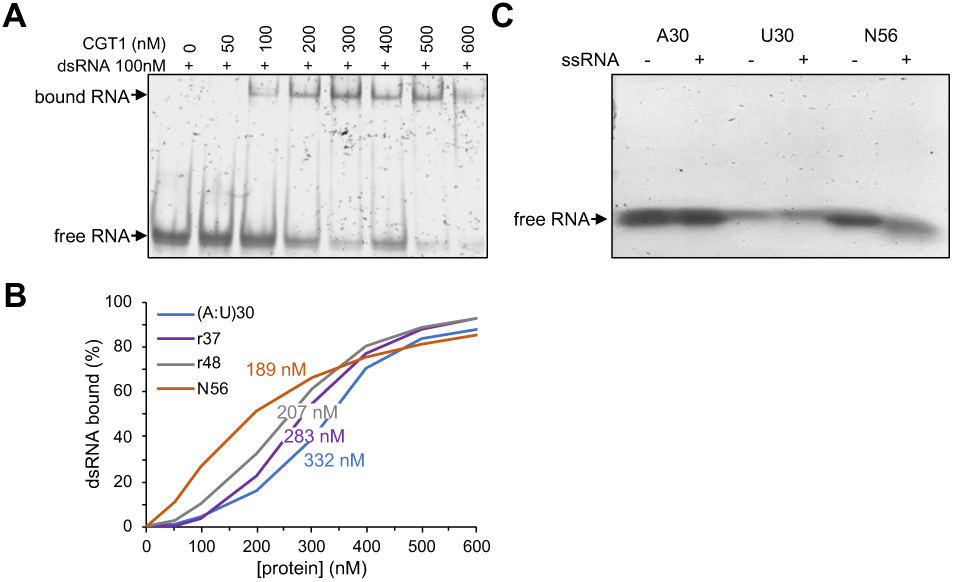
The recombinant GT1 domain binds dsRNA but not ssRNA *in vitro.* **(A)** Gel shift assay using a recombinant maltose binding protein-CGT1 fusion and a 56 bp dsRNA probe of a random sequence. **(B)** Summary of binding to different dsRNA probes and the inferred K_D_ for each probe. (A:U)_30_, 30 bp of A and U on the two strands; r37 and r48, randomized dsRNAs of the given lengths. Primary data are in Supplemental Fig. S3. **(C)** As in panel **A**, except the RNA probes were single-stranded. A30, 30 A residues; U30, 30 U residues; N56, 56 randomized nt.

Since RNJs of bacteria and archaea have been characterized to form oligomers, mostly tetramers, we analyzed Arabidopsis RNJ in this regard. Indeed, when the purified recombinant proteins were fractionated by size exclusion, all the tested proteins excluding the GT1 domain expressed without the CTD linker, formed tetrameric structures (Supplemental Fig. S4A-B). In addition, the GT1 domain alone displayed a very low dsDNA binding activity, however, the small amount eluted as a tetramer from the size exclusion column displayed high affinity binding (Supplemental Fig. S4C). These results indicated that the CTD-linker part of RNJ is required for tetramer formation, and this structure in turn is required for high binding affinity to dsDNA.

### Conserved tryptophans are key residues for *in vitro* binding ability

Transcription factor GT1 domains contain three conserved tryptophans, one in each helix, that are required for DNA binding. The RNJ GT1 domain also includes these three residues (Fig. 4A and Supplemental Fig. S1). To investigate their potential involvement in nucleic acid binding, three mutant variants of recombinant CGT1 were produced, with each of the three tryptophans mutated into alanine (Supplemental Fig. S5A). Their ability to bind the 4GT dsDNA probe was tested, with none showing any binding (Fig. 4B, Supplemental Fig. S5B-D). In parallel, ability of the W821A variant to bind dsRNA was assayed, and only weak binding was observed at the highest protein concentration tested (Fig. 4C). Therefore, each of the three conserved tryptophans of the RNJ GT1 domain is required for nucleic acid binding *in vitro*.

**Figure 4.**
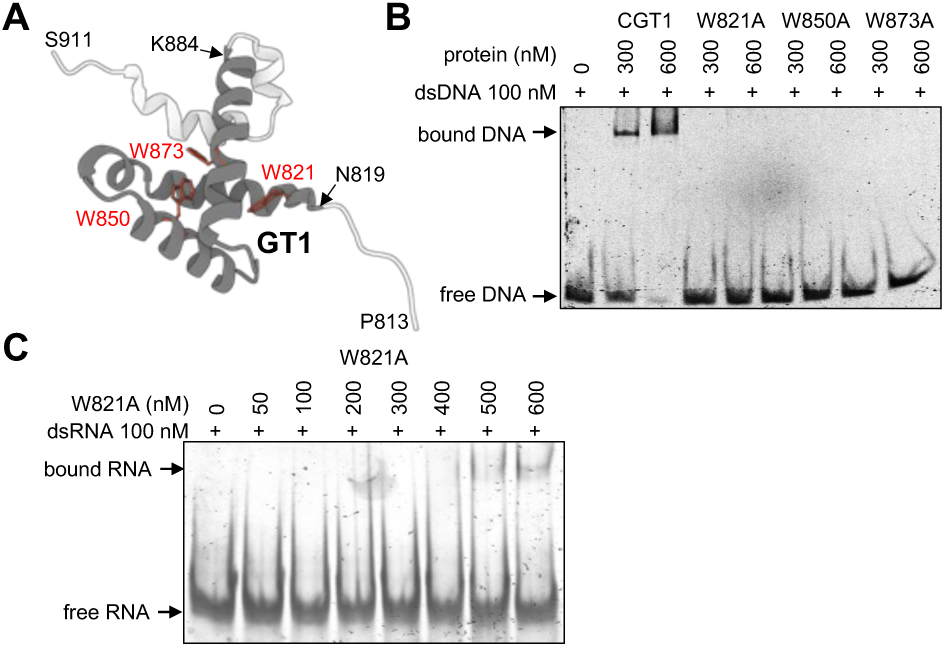
Conserved tryptophans are required for *in vitro* binding by the GT1 domain. **(A)** The GT1 domain of AtRNJ from AlphaFold structure AF-Q84W56-F1, with the three conserved Trp residues labeled in red. The orientation is rotated 180° from the structure in Fig. 1 for purposes of illustration. **(B)** A gel shift assay was carried out as described for Fig. 2A, using the 4GT probe. **(C)** Gel shift analysis was carried out as in panel **B**, except the probe was dsRNA N56 and only W821A was used.

### Two GT1-deficient RNase J variants can complement the *rnj-2* null mutant

Having characterized the binding properties of the RNJ GT1 domain *in vitro*, we wished to determine its function *in vivo*. To do so, we used complementation of the T-DNA insertion null mutant *rnj-2*, which is embryo-lethal but can be complemented by a full-length *RNJ* genomic clone including its native promoter (8). We created the three constructs shown in Figure 5A, which are driven by a 2.0 kb promoter similar to what was previously used (8), but followed by cDNAs rather than genomic sequences encoding AtRNJ. We used either the full-length version (WTc; c=control), or two versions with deficient GT1 domains, all flanked by a Flag tag. The two mutant versions were mGT, in which the three conserved tryptophans were mutated into alanines, inactivating binding, and ΔGT, a truncated version lacking the GT1 domain, which based on *in vitro* data should retain normal nuclease activity (12).

**Figure 5.**
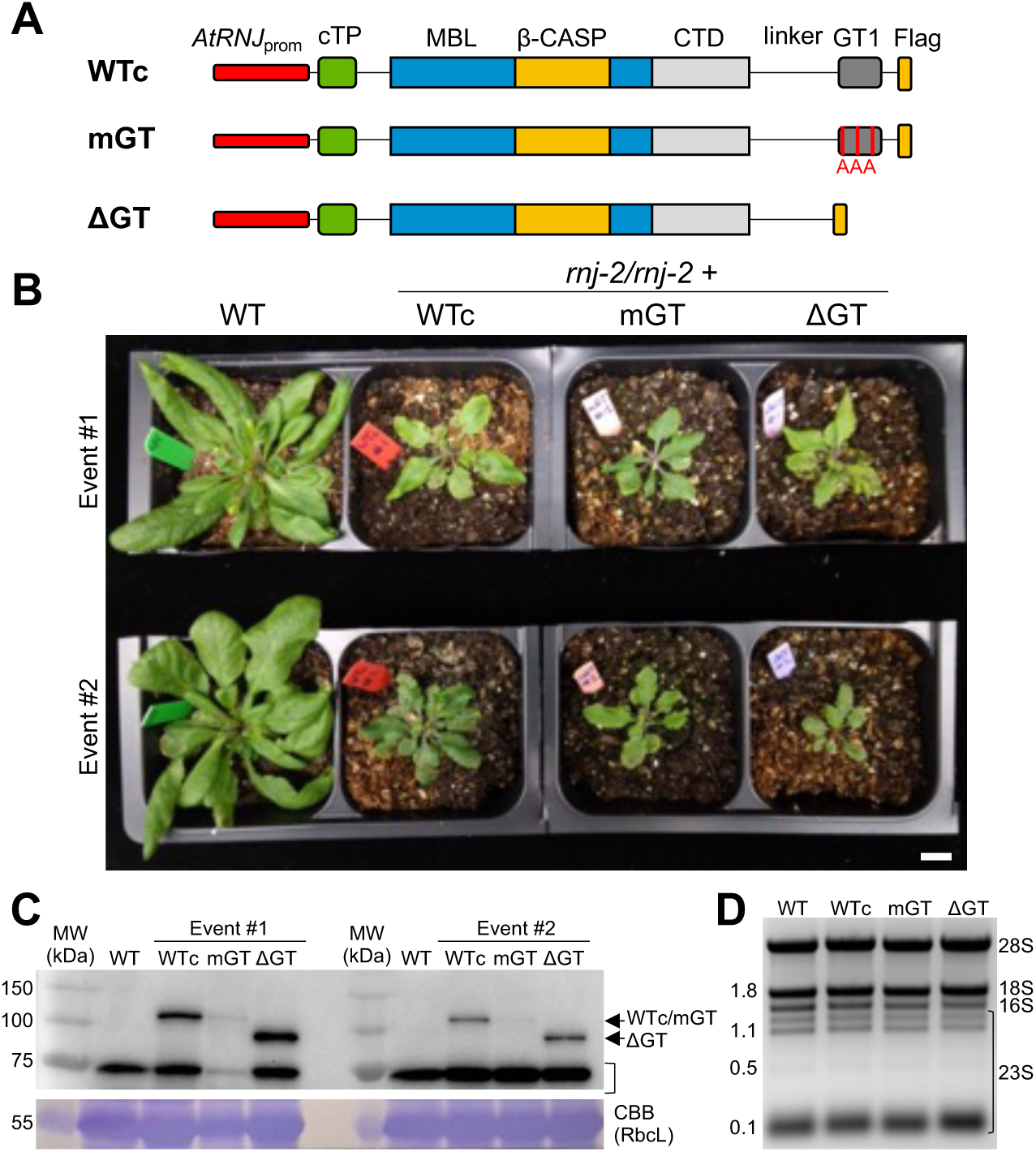
Transgenic complementation of the *rnj-2* mutant. **(A)** Constructs used to transform the heterozygous *rnj- 2* mutant. The native *RNJ* promoter was used to drive expression of AtRNJ with a WT or mutated GT1 region, flanked by a Flag epitope tag. **(B)** Untransformed WT Arabidopsis, or two representative independent events of the indicatedconstructs were grown for one month in long day conditions at 22°C (scale bar = 1 cm). Each of the transgenic plants is homozygous both for the transgene and the *rnj-2* mutation as verified by genotyping and genetic crosses. **(C)** Protein analysis of the lines shown in Panel **B**. Top, immunoblot with anti-Flag antibody showing accumulation of the three transgenic forms of RNJ. The band of ∼75 kD indicated by a bracket is present in all lines and is an unknown crossreacting protein. Lower, Coomassie-stained gel cropped to show the region of the Rubisco large subunit. **(D)** Ethidium bromide-stained gel of total RNA from the indicated genotypes. Sizes in kb are shown to the left of the gel, and major rRNA species are indicated on the right, with 28S/18S comprising mainly cytosolic rRNA and 16S and 23S being chloroplastic.

For complementation, we transformed heterozygous *rnj-2*/+ plants and screened for progeny that contained the transgene and were homozygous for the *rnj-2* T-DNA insertion. The presence of such progeny would indicate that complementation had occurred, and indeed multiple independent transformation events for each construct were identified. Based on Mendelian heritability indicating a single insertion site and phenotypic consistency suggesting a lack of other segregating mutations, two events were selected for each construct. The two events for each construct behaved virtually identically in all assays performed.

In terms of whole plant phenotype and compared to the untransformed wild-type, the complemented lines were smaller and contained less pigment, however, there was no substantial difference between WTc, mGT and ΔGT, meaning that the GT1 mutations did not affect plant growth at a macro level (Fig. 5B, Supplemental Fig. S6). To ensure that the original T-DNA mutation was not affecting plant growth we examined expression of At5g63410, which encodes an LRR protein and is 2.1 kb upstream of the native *RNJ* locus on the opposite strand. Quantitative RT-PCR showed that transcript accumulation for At5g63410 was greater or equivalent in T-DNA-containing lines (Supplemental Fig. S7A). Therefore, the three lines appeared to differ only in the form of RNJ being expressed.

Transgenic protein accumulation was evaluated by immunodetection of the Flag epitope and proteomics. The transgenic protein was clearly detectable in the three transformants, being more abundant in ΔGT and lower in mGT, compared to WTc (Fig. 5C), a result consistent with quantitative proteomic data (Supplemental Fig. S7B and Supplemental Data S1A). It was not possible to compare the level of transgenic RNJ to RNJ abundance in WT plants, due to lack of a suitable antibody. As a proxy measure, we assessed RNA abundance using RT-qPCR (Supplemental Fig. S7A). The results showed similar expression of the three transgenes relative to the actin control, suggesting that the lower level of the mGT protein relative to WTc and ΔGT is not related to RNA abundance. Transgene RNA abundance was about five-fold higher than that of the native *RNJ* gene, suggesting that RNA abundance was not a limitation to complementation, and possibly that the RNJ protein level is higher in the transgenic lines than in WT plants.

Chloroplast gene expression mutants often affect rRNA abundance and/or maturation through direct or pleiotropic effects (33). These examples include various chloroplast ribonuclease mutants, for example those affecting polynucleotide phosphorylase (PNPase), RNase II, mini- RNase III and YbeY (34–37). Similarly, plants depleted for RNJ by virus-induced gene silencing show reduced plastidial rRNA accumulation, probably as a pleiotropic effect (10). The complemented lines studied here, however, showed a normal rRNA pattern after total RNA separation in a 1.2% formaldehyde-agarose gel, implying no major change in the rRNA metabolism (Fig. 5D). We conclude that expression of RNJ with a mutated or truncated GT1 domain can complement the embryo lethality of homozygous *rnj-2/rnj-2* mutants and support plant growth as efficiently as complementation through expression of an unmutated protein. Notwithstanding, to avoid any pleiotropic effects of slower growth (Fig. 5B), we used WTc rather than untransformed plants as a control in the subsequent analyses.

### GT1-deficient lines accumulate dsRNA

The molecular phenotypes of the complemented lines were further characterized to uncover the *in vivo* function of the GT1 domain. As shown above (Fig. 3), the GT1 domain binds dsRNA *in vitro*, therefore we sought to determine whether this was also the case *in vivo*. To do so at a gross level, the transgenic proteins were immunoprecipitated using the Flag epitope, and the presence of co-immunoprecipitated dsRNA was tested using dot blot analysis with an anti-dsRNA antibody. Although dsRNA was readily detected in the IP fraction from the WTc line, little to no signal was observed for the GT1-deficient lines (Fig. 6A). This result suggests that *in vivo*, there is dsRNA present that can be bound by the GT1 domain, presumably derived from the chloroplast where RNJ is localized. Proteins lacking the GT1 domain, or with the mGT mutations, were largely ineffective in recovering dsRNA.

**Figure 6.**
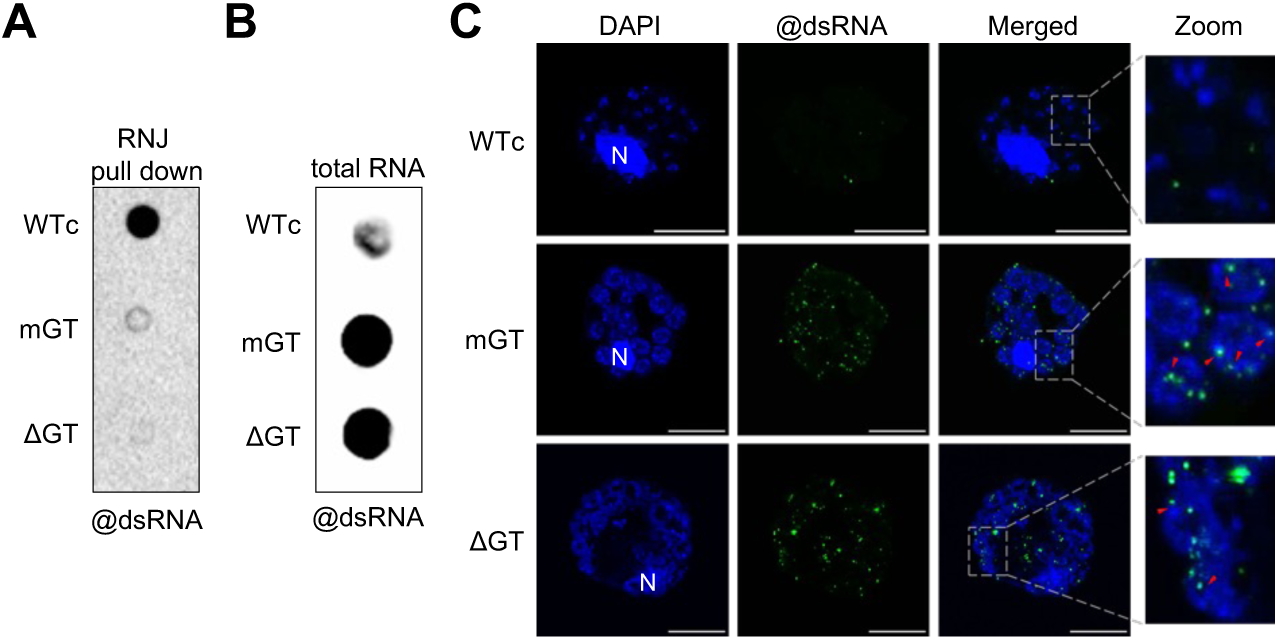
GT1-deficient lines accumulate dsRNA. **(A)** The anti-Flag antibody was used to immunoprecipitate RNase J from the indicated genotypes, then RNA was extracted and applied to a dot blot, which was probed with the J2 antibody against dsRNA. **(B)** Total RNA was extracted from the indicated genotypes, and 1 µg of each was applied to a dot blot, which was probed with the J2 antibody against dsRNA. **(C)** Immunolocalization of dsRNA (in green) using the anti-dsRNA monoclonal antibody J2 in protoplasts from the WTc, mGT, and ΔGT transgenic lines. DNA from nucleoids (in blue) was stained with DAPI. On the right, a magnified view of the protoplast is shown. The red arrows indicate regions with no overlap between green and blue signals. The scale bar for the first three columns represents 10 µm and N in the first column indicates the nucleus.

Because the results in Figure 6A could result from different levels of dsRNA in the plants, we checked overall dsRNA accumulation using the same antibody. Compared to WTc, the two GT1-deficient lines had much stronger signals, corresponding to higher accumulation of dsRNA (Fig. 6B). Thus, the lack of immunoprecipitated dsRNA by mGT and ΔGT does not reflect an absence of dsRNA from those cells, in fact dsRNA accumulation is possibly higher because those versions of RNJ are unable to bind it.

To confirm the presence of dsRNA detected in Figures 6A-B by an independent method, the same anti-dsRNA antibody was used for immunolocalization in DAPI-stained protoplasts of the complemented lines. DAPI was used both to stain the nucleus, and also the aggregations of chloroplast DNA encompassed within their nucleoids. While DAPI staining results were similar in all lines (left column), dsRNA staining was much more evident in mGT and ΔGT (speckles in @dsRNA column), consistent with the results in Figure 6B. Where dsRNA reactivity occurred, it formed granule-like foci that localized in close vicinity of chloroplast DNA (Fig. 6C). Because the nucleoid is a dense and somewhat diffuse complex containing not just DNA but also RNA and various proteins involved in gene expression (16), dsRNA foci appearing to be close to DAPI foci may in fact be within the nucleoid. Taken together, our results show that *in vivo*, the RNJ GT1 domain binds the modest amount of dsRNA that accumulates, whereas when no functional GT1 domain is present, higher levels of dsRNA accumulate and may be associated with chloroplast nucleoids. This raises the possibility that the GT1 domain helps RNJ to degrade these dsRNA by binding them and bringing them closer to the enzyme’s active site.

### GT1-deficient lines show modest changes in chloroplast mRNA abundance but not maturation and no significant changes in the accumulation of chloroplast-encoded proteins

#### Large-scale transcript and protein analysis

While the accumulation of dsRNA was a significant consequence of mutating the GT1 domain, it was possible that there would be additional effects on chloroplast RNA maturation in the complemented lines, since RNJ is also known to be involved in 5’ end maturation (11, 38). We therefore sequenced rRNA-depleted total RNA from the three complemented lines. We initially used library results to verify expected proportional representation of the three DNA-containing cellular compartments (Supplemental Fig. S8A). It was also used to zoom in on the *RNJ* locus to verify and measure the expression of the transgenes (Supplemental Fig. S8B, Supplemental Data S1A).

Since the three transgenic lines were phenotypically similar, we did not expect to find dramatic changes in the expression of nuclear genes between lines, and this was indeed the case. Statistical analysis identified >1,000 differentially expressed genes (DEGs) in pairwise comparisons (Supplemental Data S2A); however, the majority represented minor changes in gene expression, including when we selected specific gene sets related to chloroplast function. A large proportion of DEGs were shared when comparing one of the GT1-deficient lines to the WTc (Supplemental Fig S9). We used several enrichment analyses to look for any patterns, including GO, KEGG, and MapMan (Supplemental Data S2B-D). Our general conclusion is that the enriched terms identified from DEGs are similar between the mGT-WTc and ΔGT-WTc comparisons (Supplemental Table S3), reinforcing the idea that mGT and ΔGT are more similar to each other than they are to WTc. Additionally, a mGT-WTc proteomic analysis confirmed the minor consequences of GT1 deficiency on gene expression of chloroplast-encoded proteins (Supplemental Data S1B), and that no significant changes were found in splicing or RNA editing (Supplemental Data S2E-F and Supplemental Fig. S10). Taken together, we concluded that the major effect of RNJ mutations was likely to be found in more detailed analysis of the chloroplast transcriptome.

#### Chloroplast RNA differentially expressed regions

We next examined the chloroplast transcriptome in more detail. Our goal was to assess any evidence for alterations in 5’ end maturation as a result of inactivating the GT1 domain, as well as more generally find differentially expressed regions (DERs). Previous work showed that RNJ is involved in 5’ end maturation of *psbA*, *atpH*, *rbcL*, and *petB* (11), as predicted by the PPR protein-mediated RNA processing model (39). To take a systematic approach to query read coverage across the plastome, for each pairwise comparison and strand, we used the “DiffSegR” package (25) to segment the log2 ratio of coverage into regions without using existing gene annotation, generating a list of differentially expressed regions DERs between genotypes, which are displayed by strand and comparison in Figure 7A. The mGT-WTc and ΔGT-WTc comparisons showed 115 and 98 DERs respectively, with 76 and 72 being higher in WTc than the GT1-deficient lines (red color). When the two GT1-deficient lines were compared, however, only 30 DERs were identified (Fig. 7A-B, Supplemental Data S2G, Supplemental Fig. S11). The patterns of DERs identified by comparing GT1-deficient lines to WTc were highly similar at a coarse visual level, consistent with effects on the chloroplast transcriptome being substantial and conserved in the two GT1-deficient genotypes.

**Figure 7.**
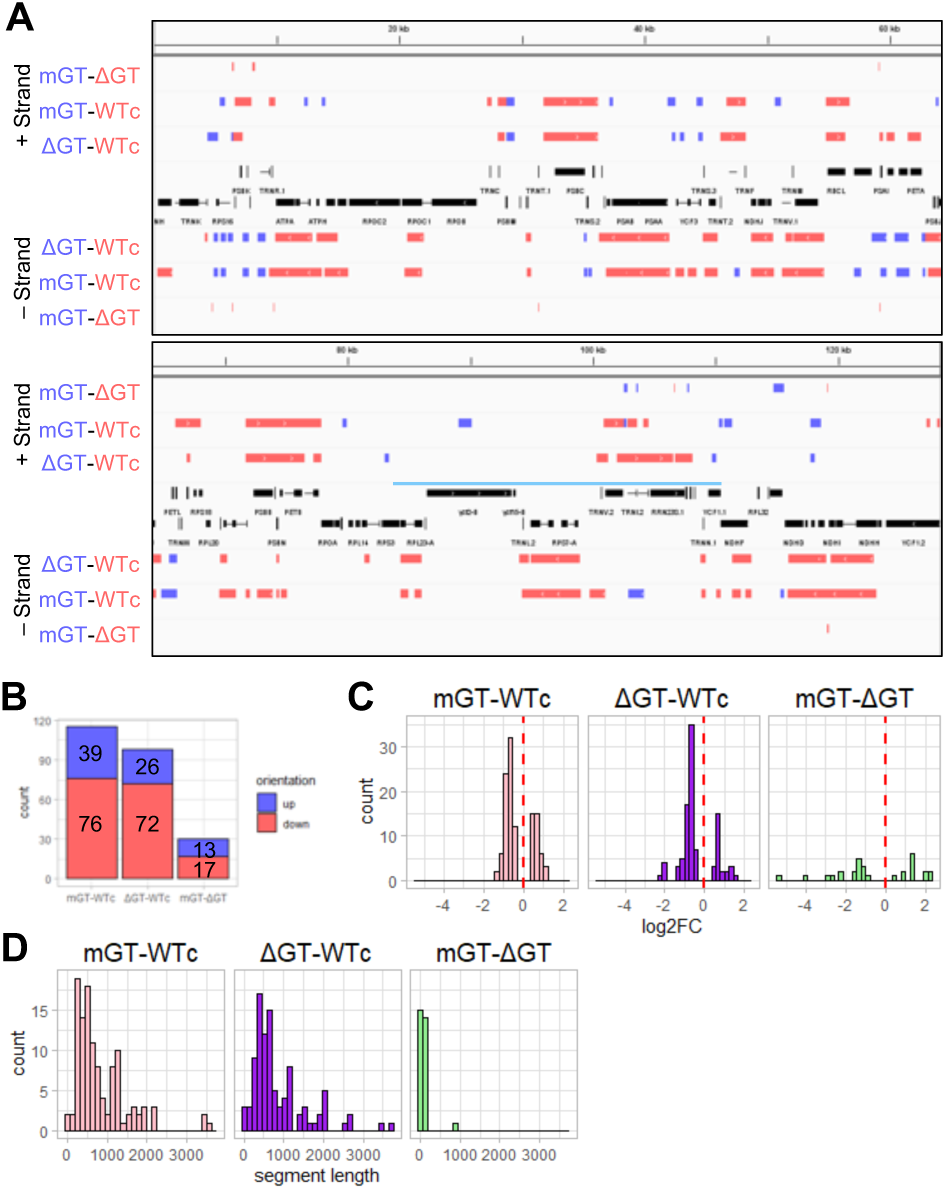
Pairwise comparisons of differential accumulation of transcripts in complemented lines. **(A)** Localization of DERs by strand and pairwise comparison, using visualization on IGV of chloroplast genome coordinates 1- 128,216, thereby excluding one copy of the inverted repeat. Selected genes are annotated in black at the center. DERs are color-coded by being up-regulated (blue) or down-regulated (red). The light blue line denotes the extent of the inverted repeat copy shown. **(B)** DER count for each pairwise comparison, corresponding to the data displayed in Panel **A**. **(C)** Log2 fold change for all DERs identified in Panels **A** and **B**. The graphs combine up/down DERs. **(D)** Length distribution of DERs as displayed in Panel **C**.

To delve further into the identified DERs, we examined their amplitude, length, and location (Fig. 7C-7D). In terms of amplitude, most changes were within a twofold range, and the patterns were similar between mGT and ΔGT vs. WTc. As noted above, there were few DERs between mGT and ΔGT, and for these the amplitudes were more scattered. Length analysis showed that while some DERs were short (nearly all for mGT-ΔGT), about half were 500 nt to 2 kb or larger. This suggests that differential transcript accumulation in mGT and ΔGT occurred over regions rather than in specific locations such as RNA termini.

#### 5’ end maturation

Maturation defects should generate 5’ (or 3’) extended transcript species, leading to the identification of a DER upstream (or downstream for 3’ maturation) of the gene with a greater amplitude than the observed inter-genotype variations within the cognate gene. In contrast to this prediction, DERs seem to mainly accumulate within gene annotated regions, and no drastic changes were seen between a gene and its flanking regions. In other words, across individual genes or gene clusters, even when several adjacent DERs span it, we found the coverage ratio to be roughly uniform along gene and flanking loci, giving no hint of terminal maturation defects (Supplemental Fig. S12). We also employed 5’ RACE to check the individual transcripts previously investigated (11). As with RNA-seq, no reproducible differences were found between genotypes (Supplemental Fig. S13), suggesting that the GT1 domain does not have a role in this process, and consistent with the demonstration that a commercial 5’ to 3’ exonuclease can fulfill this role *in vitro* (39). We conclude that mutation of the GT1 domain causes quantitative changes in the chloroplast transcriptome, but not to the maturation of individual annotated transcript termini.

#### Summary of DER patterns

DER distribution was analyzed with respect to chloroplast chromosome annotation as exons, introns, the antisense strand of exons and introns, and intergenic regions, with the exception of tRNAs and rRNAs which are depleted during library preparation. To do so, we used DER coordinates to calculate coverage density, meaning the proportion of the delineated space (exon, intron, etc.) that is covered by DERs. The results for this are shown in Figures 8A and 8B, where the Y-axis shows that proportion, and the X-axis shows the amalgamated and normalized feature of interest, e.g. all exons regardless of actual length are shown as an arbitrary length of 1000.

**Figure 8.**
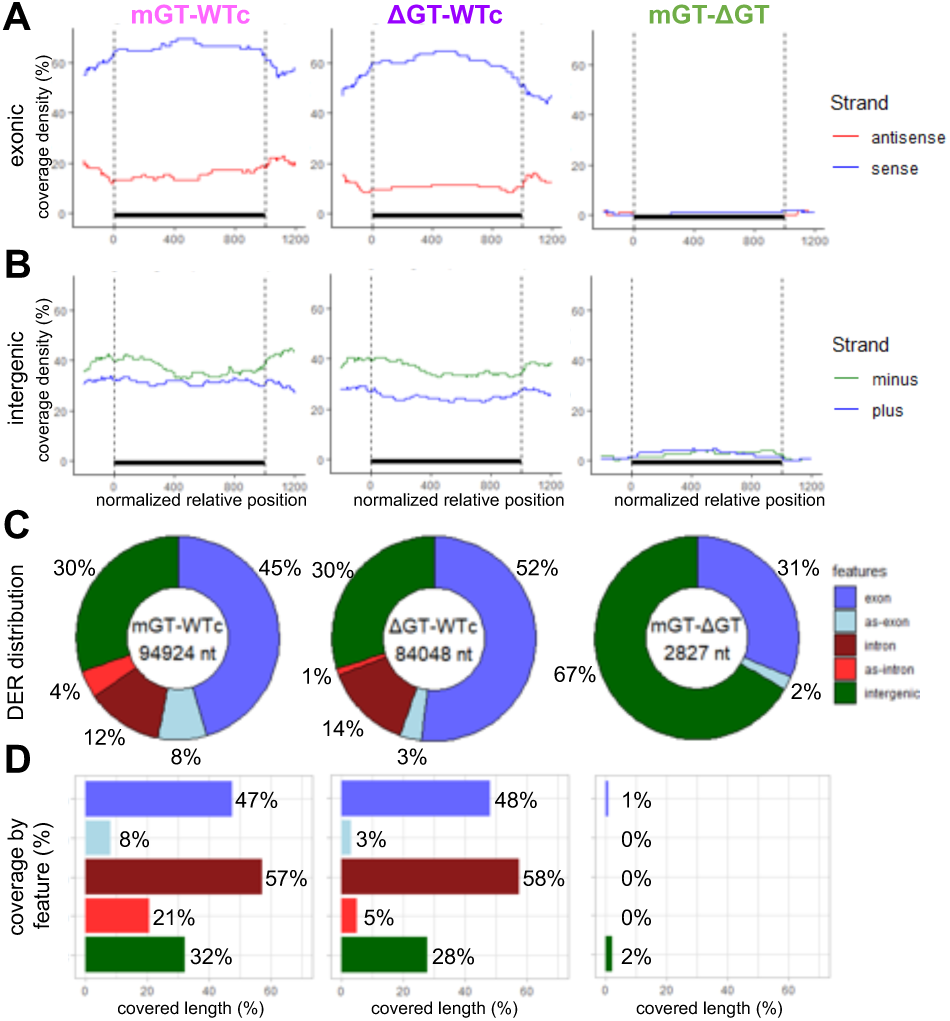
Distribution of DERs in exonic and intergenic regions. **(A)** Average DER coverage over protein-coding exons (n=105) for each pairwise comparison. Blue lines correspond to the sense strand orientation; red to the antisense orientation. **(B)** Average DER coverage over intergenic regions (n=128) for each pairwise comparison; intergenic regions were defined as described in Materials and Methods. Blue lines correspond to the coverage on the plus strand; green to the minus strand. **(C)** DER coverage distribution by genome feature for each pairwise comparison. The total number of bases in the DERs for a given pair is shown at the center of the diagram. Genomic features are color-coded as exon (blue), antisense-exon (light blue), intron (brown), antisense-intron (red), or intergenic (green). **(D)** Proportion of each type of genomic feature covered by DERs for each pairwise comparison. Genomic features are color-coded as in panel C.

This analysis was quite striking when WTc was compared to either of the GT1-deficient lines, where on average about 2/3 of exon space is differentially expressed when those two genotypes are compared. In contrast, there was low DER coverage on the exon antisense strand, and basically no DER coverage when the two GT1-deficient lines were compared. Combining this analysis with the data shown in Figure 7A, it is obvious that the DERs are mainly composed of underexpressed regions in the GT1-deficient lines. We can therefore conclude that the sense, but not antisense strand of exons tends to decrease when the GT1 domain is not present. When intergenic regions were plotted collectively, DER coverage was 30-40% but not strand-preferential.

We also summarized DER distribution by the collective lengths of DERs across features (Fig. 8C) and as a proportion of the collective length of each feature over the plastome (Fig. 8D). In the center of Figure 8C the sum of all DERs lengths can be seen: this totals about 95 kb and 84 kb in the comparisons of mGT and ΔGT with WTc, respectively, whereas DiffSegR only identified 2.8 kb of DERs between the two GT1-deficient lines. DERs in mGT and ΔGT correspond to 31% and 27% of the two strands of chloroplast chromosome, meaning the chloroplast transcriptome is widely affected. Figure 8C also shows that 75% and 82% of the DERs are either intergenic or on the exonic sense strand. When this coverage is proportionalized to the summed length of these features across the genome (Fig. 8D), the exon sense strand is about half covered by DERs and intergenic regions around 30%. Introns are more than half covered, although they only represent 10-15% of DERs.

Taken together, these coverage analyses suggest an overall transcript accumulation variation that impacts the exon sense strand, introns, and intergenic regions, but do not support a role for the GT1 domain in RNA end maturation. These quantitative changes do not result in major changes in chloroplast protein abundance as assessed by proteomics (Supplemental Data S1B). This is probably because chloroplast mRNAs are present in excess in most cases (see Discussion).

### Differentially accumulating dsRNA covers a large proportion of non-coding regions

We noted above that mGT and ΔGT accumulate dsRNA (Fig. 6B), and with the techniques used the RNA-Seq reads would be derived from RNA whether single- or double-stranded, therefore the changed accumulation in mGT/ΔGT could be from either or both forms. For this reason, we next prepared sequencing libraries specifically from dsRNA.

To further characterize this dsRNA population, it was immunoprecipitated using the anti-dsRNA antibody, sequenced, and mapped similarly to total RNA, except that we focused exclusively on the chloroplast reads. Compared to the total RNA coverage, DiffSegR-generated DER profiles were markedly different (Supplemental Fig. S14, Supplemental Data S2H). First, DERs map across the chromosome on both strands, except in the rRNA and tRNA-rich inverted repeat where transcripts should be depleted (Fig. 9A). Second, rather than a mix of up and down regulation, changes are nearly exclusively increases in the GT1- deficient lines (Fig. 9B). Specifically, only 1 of 295 DERs in mGT represents down regulation, and only 14 of 216 in ΔGT. Comparison of mGT and ΔGT yielded 55 DERs, all of which represented increased accumulation in mGT (see Discussion). The amplitude of DERs trended slightly higher than what was observed with total RNA (compare Fig. 9C and 7C), whereas the lengths trended shorter (compare Fig. 7D and 9D). This profusion of likely short and substantially up-regulated regions is consistent with our previous observation showing the association between dsRNA accumulation and removal or inactivation of the GT1 domain (Fig. 6B).

**Figure 9.**
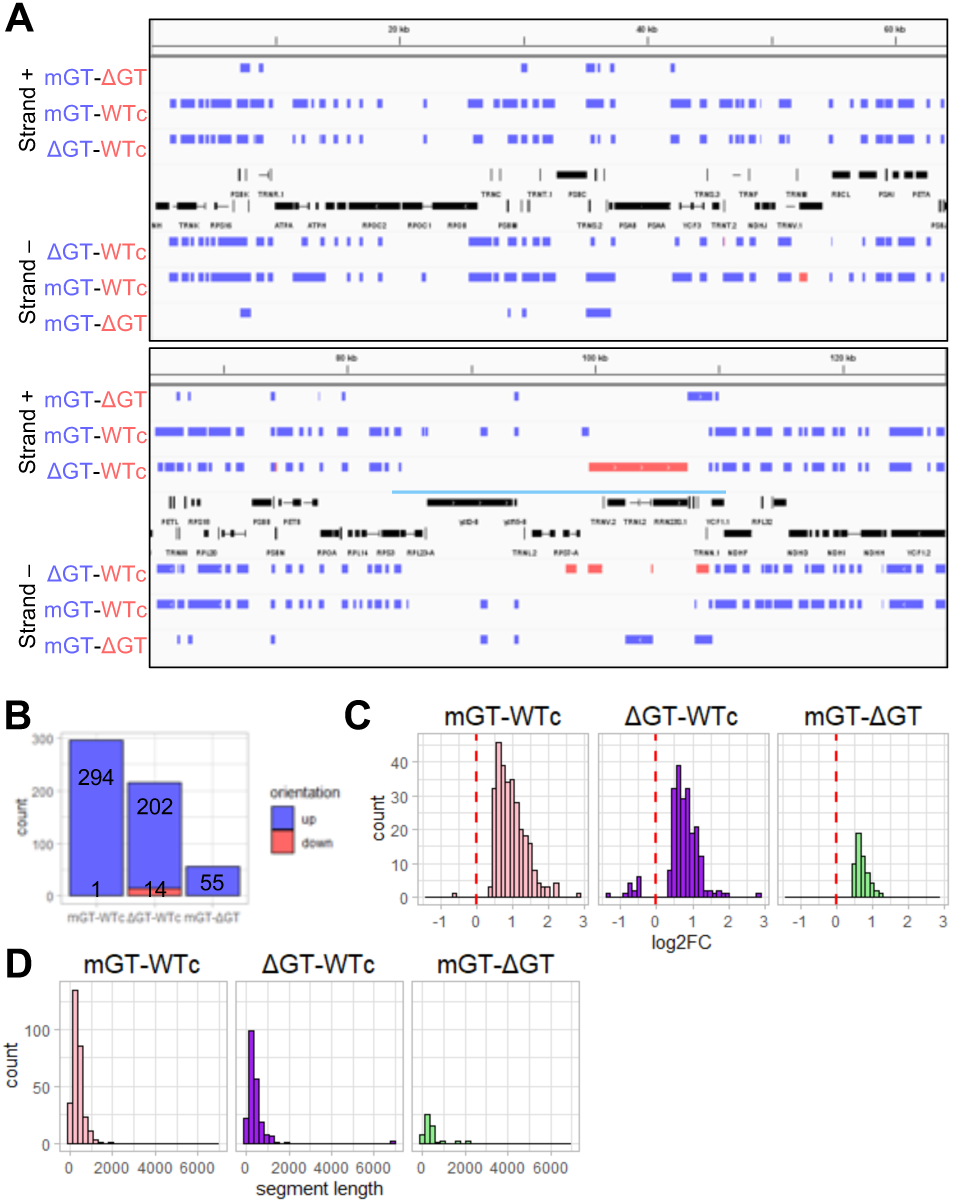
Differential accumulation of dsRNA in pairwise comparisons. **(A)** Localization of DERs derived from RNA-seq analysis of immunoprecipitated dsRNA, as described in the legend to Fig. 7A. **(B)** The number of DERs in each pairwise comparison, as in Fig. 7B. **(C)** The distribution of log2 fold changes in DERs, as in Fig. 7C. **(D)** Length distribution of DERs as in Fig. 7D.

DER coverage density was analyzed as for total RNA, where a rather different picture emerged. We found that coverage density tended to decrease over protein-coding exons, but to increase within intergenic regions with no discrepancy between sense and antisense coverage (Fig. 10A-B). About 43% of 110 kb and 100 kb covered by DERs in mGT and ΔGT, respectively, are located in intergenic regions and span about half of the total intergenic regions. About half of the intron and as-intron length is also covered (Fig. 10C-D). Together, these analyses reveal that in the absence of a functional GT1 domain, dsRNA accumulates from broadly distributed loci, primarily from intronic and intergenic regions. This best supports a generalist rather than sequence-specific function of RNase J in RNA quality control.

**Figure 10.**
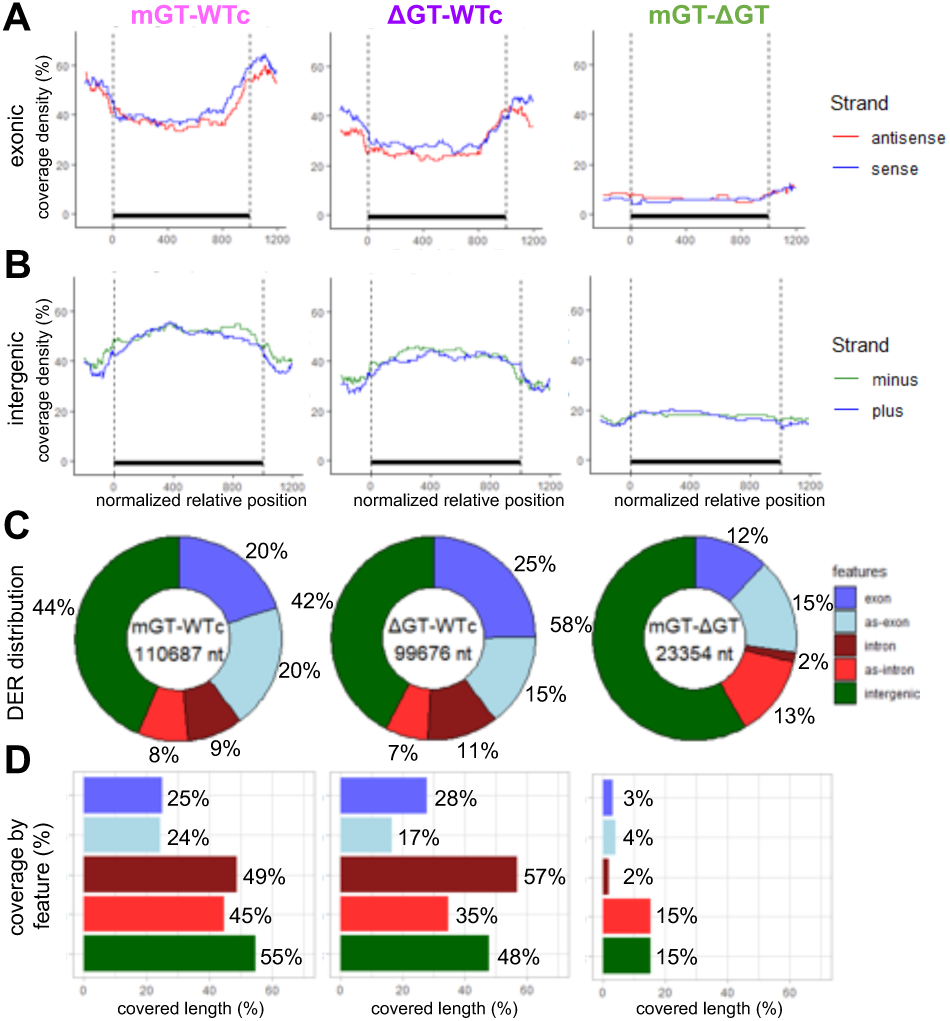
Accumulation of dsRNA by genomic region in pairwise comparisons. **(A)** Average DER coverage over protein-coding exons (n=105) for each pairwise comparison, as described for Fig. 8A. **(B)** Average DER coverage over intergenic regions (n=128) for each pairwise comparison, as in Fig. 8B. **(C)** DER coverage distribution by genome feature for each pairwise comparison, as in Fig. 8C. **(D)** Proportion of each type of genomic feature covered by DERs for each pairwise comparison, as described in Fig. 8D.

### Accumulated dsRNAs are likely to be present as sense-antisense duplexes

Double-stranded RNA can be formed by a secondary structure (intramolecular or *cis* pairing), or by hybridization of two reverse complementary molecules (sense-antisense duplexes or *trans* pairing). In plants where RNase J was silenced, asRNA accumulates and impedes with the translation by forming such duplexes (10). To attempt to differentiate *cis* vs. *trans* base pairing for the immunoprecipitated dsRNA accumulating in mGT and ΔGT, an additional coverage analysis was conducted on the dsRIP-Seq data. This was to segment within each genotype and to compare plus over minus strand coverage. In principle, immunoprecipitation of *trans*-dsRNA should lead to a symmetrical signal on both strands, whereas a captured *cis* structure would generate a signal biased toward one single strand.

Identification of symmetrical dsRNA DERs revealed 68 unique *trans*-dsRNAs across the three genotypes (Fig. 11A, Supplemental Data S2I). Although the number of unique species is similar, mGT and ΔGT share most of them and those identified in WTc seem more specific to this genotype (Fig. 11B). Arabidopsis chloroplasts are already known to accumulate a certain level of symmetric chloroplast transcripts: for example, a previous study reported 107 non-coding chloroplast RNAs (40), most of them being antisense to known genes and thus with the potential to form *trans*- dsRNAs. Another well-characterized antisense transcript is *psbN*, which is antisense to the *psbB* gene cluster (41) Coordinates of the 68 putative *trans-*dsRNAs were compared with the 107 ncRNA locations, and we found that 26 *trans*-dsRNAs overlapped with ncRNAs (Supplemental Fig. S15). Therefore, dsRNAs that accumulate in absence of a functional GT1 domain are a blend of known dsRNAs, some of which hyperaccumulate, and novel *trans*-dsRNAs resulting from a failure in RNA quality control.

**Figure 11.**
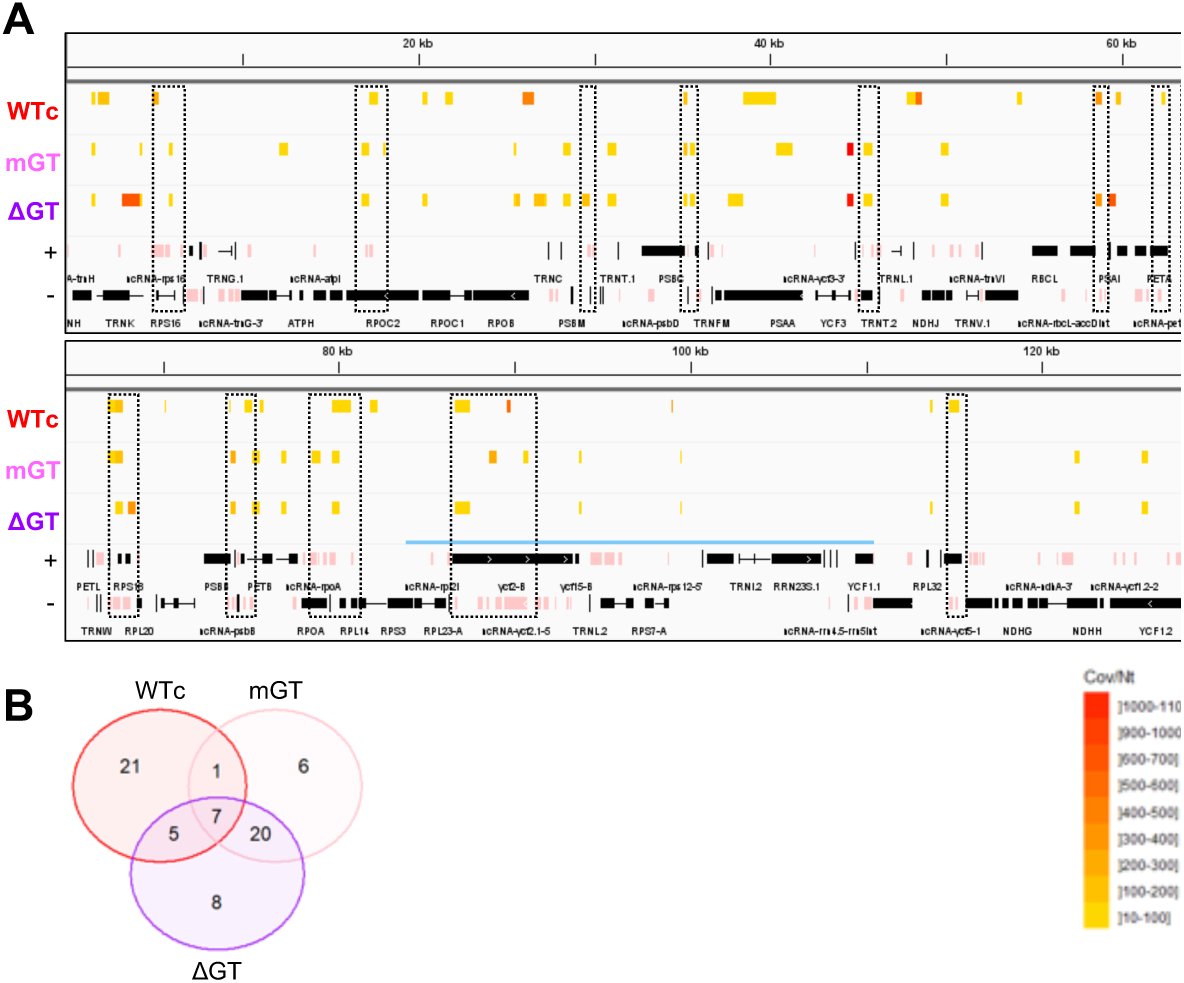
Identification of putative *trans*-dsRNAs. **(A)** Localization of the 68 unique *trans*-dsRNAs by genotype, using visualization on IGV of chloroplast genome coordinates 1-128,216, thereby excluding one copy of the inverted repeat (overlined in blue). Top to bottom, tracks represent: WTc, mGT and ΔGT, position of the putative *trans*-dsRNAs in the respective genotypes where yellow to red shades depict intensity of accumulation according to the scale at lower right; +/-, annotation of genes (black) and ncRNAs (pink) from Hotto *et al*., (2011) on the plus and minus strands. Dashed boxes highlight intersections between ncRNAs and putative *trans*-dsRNAs magnified in Fig. S15. **(B)** Venn diagram of the putative *trans*-dsRNA population.

### The GT1 domain may anchor the RNase J in chloroplast nucleoid

Our RNA-Seq results strongly suggest that the GT1 domain interacts with dsRNA *in vivo*, in keeping with the EMSA results described above. These same experiments also identified a dsDNA binding capacity, which like dsRNA binding did not appear to be sequence-specific. Given the previously reported localization of RNJ to the nucleoid (16, 17, 42–44), one obvious hypothesis is that the GT1 domain plays a role in this localization through tethering RNJ to chloroplast DNA. Since the nucleoid itself is thylakoid-membrane-associated, an approach was taken that would partition proteins between a soluble phase and a membrane-containing insoluble phase that should also include nucleoid components.

In a first experiment that was minimally invasive, total proteins were separated into such soluble and insoluble fractions, and the partitioning of Flag-tagged RNJ was evaluated by immunoblot. This analysis was restricted to WTc and ΔGT due to the low expression level of RNJ in mGT (Fig. 5C). The results shown in Figure 12A for two events of each genotype, show different ratios for the WTc and ΔGT proteins when the soluble and insoluble fractions are compared. Specifically, the signal for WTc was 2-3X higher in the insoluble fraction, whereas the signal for ΔGT was similar between fractions. In a second approach, chloroplasts were first purified and then thylakoid and stromal fractions were isolated. Figure 12B shows the same trend as in the whole cell approach, where WTc is more abundant in the membrane fraction and ΔGT is more abundant in the stromal fraction. Thus, the two methods yielded consistent results, and support the hypothesis that the GT1 domain may anchor the protein within the nucleoid through its ability to bind non-specific sequence of dsDNA. Because the partitioning was not absolute for either protein, it is possible that RNJ is present in two populations in both transgenic lines, meaning that its apparent solubility is not solely determined by the GT domain.

**Figure 12.**
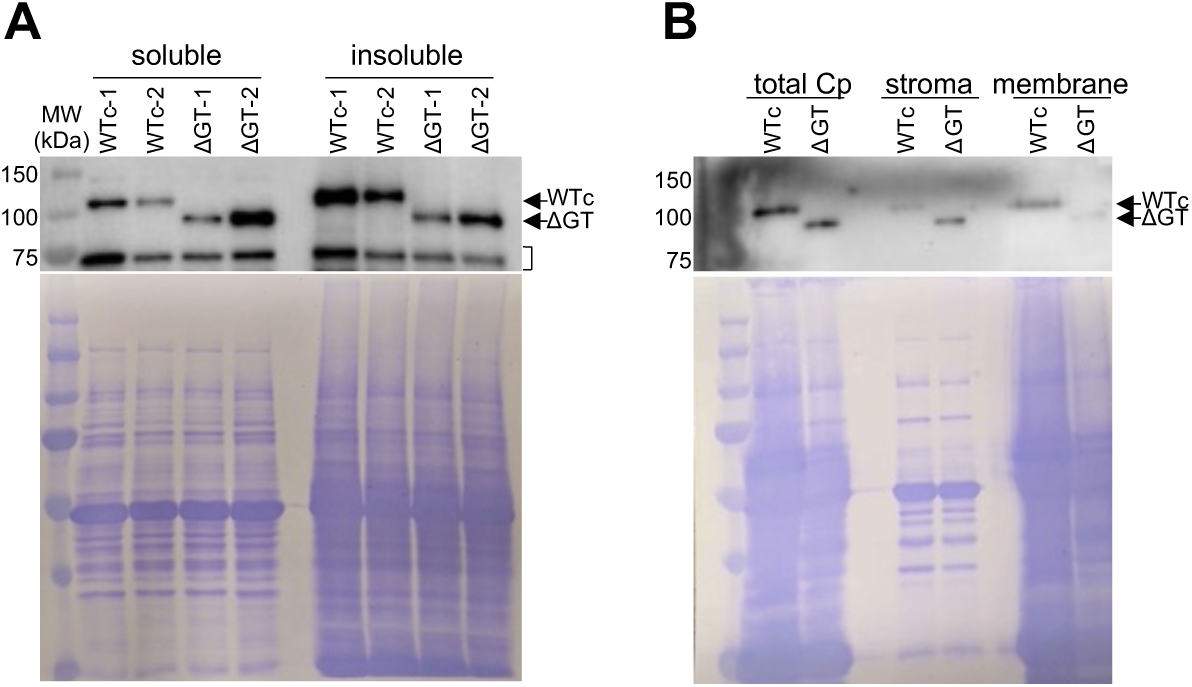
The GT1 domain may affect RNJ solubility.**(A)** Total leaf protein analysis of both events of WTc and ΔGT. Top, immunoblot with anti-Flag antibody F3165. The band of ∼75 kD indicated by a bracket is an unknown cross-reacting protein. Lower, Coomassie-stained gel. **(B)** Total chloroplast, stromal and membrane fractions from intact isolated chloroplasts.

## DISCUSSION

### The unique presence of a GT1 domain in plant RNase J

While RNase J is found in all domains of life, only in land plants have forms been described that include a GT1 domain. This domain is widely found in a subset of trihelix transcription factors (15), where the DNA-binding domain is hypothesized to be derived from transposable elements (45). We speculate that the GT1 domain in RNase J was similarly derived after the divergence of plants from organisms whose RNase J’s do not contain GT1 domains, and that its maintenance either conferred a functional advantage or simply persisted as a neutral trait. The work presented here was designed to assess what that functional advantage might be, through complementation of an *rnj* mutant with transgenes expressing forms of RNase J either lacking the GT1 domain or containing it in a binding-disabled form.

### AtRNJ appears to bind both double-stranded DNA and RNA without sequence specificity

The literature on GT-1 transcription factors has naturally focused on their DNA-binding properties, and our working hypothesis had been that the RNase J GT1 domain would similarly confer dsDNA binding ability to RNJ, most likely with a sequence preference related to that of the canonical factors. This had led us to speculate that RNJ might be targeted to sites on the chloroplast chromosome with sequence resemblance to GT elements, and potentially act on transcripts synthesized from those regions. At the same time, the global effects on the chloroplast transcriptome of RNJ depletion (10) would argue against such a model.

Our *in vitro* binding studies of recombinant RNJ and derivatives lent clarity to these alternative models. Most important was a lack of sequence specificity, with the main parameter affecting DNA binding being the length of the probe (Supplemental Fig. S2). Stronger binding to a longer probe is consistent with earlier studies of the pea GT-1 factor, which bound *in vitro* to probes containing three or four tandem copies of the *RBCS* GT element, but not to probes of one or two copies (32). While the degeneracy of plant GT-1 binding sites is documented (46), our results for RNase J suggest that in a chloroplast chromosome targeting model, RNase J would be bound stochastically to cpDNA. Important caveats are that we did not sample sequences exhaustively, that we assayed the GT1 domain outside the context of the full-length protein and in defined buffers optimized for *in vitro* experiments, and that DNA accessibility and structure vary across the chloroplast chromosome and are likely to affect binding sites and frequencies. Sequencing of DNA bound *in vivo* to RNase J might reveal certain preferred sites for binding, if these in fact exist.

A more unexpected result was that the RNase J GT1 domain bound dsRNA *in vitro* with a high affinity (Fig. 3), a result that was strongly supported by the dependence on a functional GT1 domain for immunoprecipitation of dsRNA (Fig. 6A). While to our knowledge the dsRNA binding ability of GT-1 transcription factors had not been previously tested, related experiments had been performed for the human HMG domain transcription factor Sox2. Similar to our results for RNase J, Sox2 binds dsRNA *in vitro* in a non-sequence-specific manner and this behavior has been ascribed to the flexibility of its binding domain (47). Sox2 also can be associated with several lncRNAs (e.g. 48) just as RNase J was identified in a pulldown experiment with the plant lncRNA lncCOBRA1 (49). These results raise the possibility that certain trihelix transcription factors may have additional interactions with the transcriptome, something not well documented in plants yet exemplified already by the dual functions of the rice bZIP transcription factor APIP5 (50).

### The GT1 domain is not required for complementation of embryo lethality

We were able to complement the embryo lethality of *rnj* mutant with constructs expressing RNase J with or without a functional GT1 domain, showing that the domain is dispensable for plant growth under laboratory conditions. We observed that growth phenotypes of the three transgenic lines were indistinguishable (Fig. 5B), although all plants were smaller than the true WT. We were able to rule out poor transgene expression (Supplemental Fig. S7A) as an explanation, and position effects were ruled out by analysis of multiple independent transgene insertion events. This leaves two main possibilities to explain the smaller plant size in WTc vs. true WT: either the Flag tag impedes protein function in some way or conversely, RNJ is being overexpressed relative to WT and this leads to a dominant negative phenotype, albeit one with no clear effects on chloroplast gene expression (in WTc). We attempt to avoid the latter issue by using the native promoter, however this does not ensure a precise replication of WT protein expression and indeed, transcript levels were higher for all of the transgenes than they were for the native transcript in the true WT (Supplemental Fig. S7A). A weaker promoter could potentially promote transgenic expression that more closely matched the true WT. Any effect of the Flag tag could in principle be tested by expression of transgenes without the tag or with a different tag.

Two other studies have reported successful complementation of an *rnj* mutant. In Arabidopsis, it was shown that a genomic clone, presumably lacking an epitope tag, could complement embryo lethality (8), but the adult plant growth phenotypes were not reported and it is possible that complete recovery of growth was not observed. In rice full complementation was observed using a cDNA clone driven by the actin promoter and also presumably lacking an epitope tag, however the mutant that was complemented was a point mutant with a revertible early-stage mild chlorotic phenotype (9). Whether the precise abundance of RNase J is a critical factor for normal plant growth may be an interesting topic for future study.

The fact that Arabidopsis RNase J does not require a GT1 domain to support embryogenesis and plant growth suggests that its function is regulatory in nature. Indeed, the absence of a GT1 domain in the Chlorophyte *Chlamydomonas reinhardtii* (51)as well as in bacterial and Archaeal RNase J, points to an adaptive function in flowering plants. Genomic structural differences that could rationalize a regulatory role in flowering plants are discussed further below.

### Removing or inactivating the GT1 domain increases dsRNA accumulation and may decrease mRNA sense strand abundance

Given that VIGS of RNase J led to accumulation of dsRNA in tobacco (10), and that our *in vitro* experiments demonstrated a capacity of the GT1 domain for dsRNA binding, investigation of dsRNA abundance and sequence in our transgenic lines was important to ascertain. When an anti-Flag antibody was used to immunoprecipitate the different forms of RNase J and the pellet was evaluated for the presence of dsRNA, only WTc presented a signal (Fig. 6A). This suggests that dsRNA is tightly bound to RNase J in WTc, and that the GT1 domain is required for this binding. When total RNA was assayed for the presence of dsRNA, its abundance was much higher in mGT and ΔGT than in WTc (Fig. 6B). This result suggests that binding of dsRNA to RNase J is linked to its degradation, and this process requires a functional GT1 domain to be highly efficient. Our results do not allow a comparison of dsRNA abundance between our lines and the VIGS plants created by Sharwood *et al*., however we speculate that VIGS induced higher dsRNA accumulation given the dramatic phenotypic effect that was observed.

Transcriptome analysis of dsRNA showed that much of the genome was present in this form and of much higher abundance in mGT and ΔGT than in WTc (Fig. 9-10). This finding is in keeping with the apparent lack of sequence specificity of GT1 binding *in vitro*. Nonetheless, coverage density was lower in exonic regions than in intergenic regions, which might explain in part why plant growth was possible in spite of the presence of asRNA that might be expected to impede translation. Whether a given asRNA would actually inhibit translation would depend on a number of factors including its length, sequence, stoichiometry, and ability to compete with *cis* folding of each strand. Some of these considerations have been investigated in the context of designing asRNAs for synthetic biology applications (52). It has also been shown that chloroplasts mRNA abundance may be well in excess of what is required for protein homeostasis (53), which would leave room for a certain degree of translation inhibition without phenotypic consequences.

Antibodies raised against dsRNA are designed to recognize fully-paired duplexes of any sequence and sufficient length (e.g. >40 bp), and not to recognize ssRNA or RNA-DNA hybrids (54, 55). They should also react poorly with partial duplexes such as those found in rRNA and tRNA. Nevertheless, we cannot exclude that some of the molecules identified in our study represent *cis* interactions such as intron domains, rather than *trans* interactions of sense- antisense strand pairing. We used the presence of symmetrical sense and antisense strands to identify putative *trans*-dsRNAs (Fig. 11), however that method is correlative rather than definitive. Our overall conclusion that mGT and ΔGT accumulate abnormally high levels of chloroplast dsRNA, however, is well supported by the range of assays used.

While pairwise transcriptome comparisons of WTc, mGT and ΔGT did not show any significant changes in nuclear gene expression or chloroplast protein abundance, we did note a decrease in the abundance of chloroplast coding regions using a segmentation tool and binning DERs covering coding regions (Fig. 8A). The decreases observed using this approach did not manifest in intergenic regions nor on the antisense strand (Fig. 8A-B). A more classical DEG analysis showed an opposite trend for some genes (Supplemental Fig. S9E), likely due to differences in normalization (based on chloroplast expression for DiffSegR versus total gene expression for DEG analysis) among other methodological variations.

If one accepts for the sake of discussion that GT1 mutations lead to a decrease in the abundance of exonic sequences, this would be consistent with a role in co-translational RNA degradation. RNA quality control is important during translation because transcripts may be defective due to premature termination or faulty processing and require removal from the ribosome and subsequent degradation (reviewed in 56, 57). While these pathways are well documented for nucleus-encoded mRNAs, little is known of how the chloroplast handles RNA defects during translation. Further analysis of RNase J could open a window into this putative regulatory function.

### The significance of the GT1 domain binding to both DNA and RNA

We have shown that the GT1 domain, when present in a tetramer, can bind dsDNA and dsRNA *in vitro*, raising the question of whether both of these interactions occur in the plastid, and contribute to the observed transcriptome and protein solubility phenotypes. DNA binding could act as a tether to the nucleoid, which could be important for RNA quality control soon after transcription. We speculate that dsRNA binding is important for locating duplex RNAs that need to be degraded, whether present in the nucleoid or elsewhere. Since the mutant RNase J’s bind neither DNA nor RNA, our studies cannot differentiate between the importance of interactions with the two substrates. Indeed, there are many instances of proteins that bind both DNA and RNA in nature, which can be cooperative, competitive or independent (58). Our results presented here show a slightly higher affinity for dsRNA than dsDNA (compare Fig. 3B and Supplemental Fig. S2O), suggesting that dsRNA could outcompete DNA binding to the GT1 domain in some circumstances. RNJ also has a catalytic site which is thought to bind exclusively ssRNA, although it is possible that either binding or catalysis could be influenced by what is (or is not) bound at the GT1 site. Measuring catalytic activity with or without GT1 binding substrates could illuminate this issue.

Another chloroplast-localized protein shown to bind both DNA and RNA is Whirly1, which was originally identified in maize by its interaction with the *atpF* splicing factor CRS1 (59). Co-immunoprecipitation showed non-sequence-specific interaction with chloroplast DNAs, but a higher level of specificity with RNAs. In contrast to the RNase J GT1 domain, Whirly1 bound preferentially to single-stranded RNA and DNA *in vitro*, with no interaction observed for dsRNA. It has been proposed that the DNA binding activity of Whirly1 tethers it to the thylakoid membrane and is important for nucleoid assembly (60), whereas the RNA-binding activity promotes *atpF* splicing. Other studies, however, suggest that the DNA-binding function is of minor importance (61), or that Whirly1 may be involved in diverse functions including DNA replication and retrograde signaling (reviewed in 62). These results point to the complexity of dissecting the individual roles of multifunctional proteins.

#### A model for RNJ activity

A working model for RNJ that encompasses its known properties is shown in Figure 13. We hypothesize that like Whirly1, RNase J is tethered to the nucleoid, in this case via the GT1 domain. The increase in RNase J solubility in ΔGT (Fig. 12A) supports this model, along with the identification of RNase J in the nucleoid proteome (16, 17). The chloroplast nucleoid is known to be a site of transcription and RNA processing (16), and RNase J’s documented role in 5’ end maturation could be enhanced by a nucleoid localization. The fact that the mGT and ΔGT lines did not have defective 5’ end maturation suggests that nucleoid localization is not required for this function of RNase J, and indeed the proteomic studies cited above do not propose an exclusively nucleoid localization.

**Figure 13.**
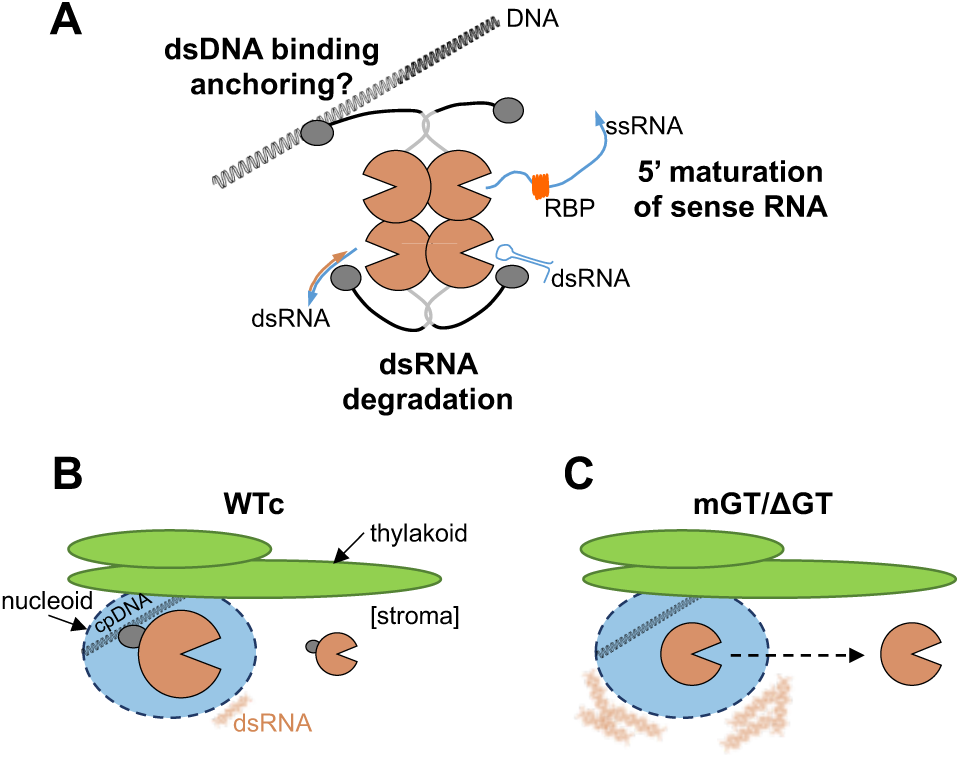
Model for RNJ functions *in vivo.* **(A)** Activities of RNJ in a WT plant. RNJ is shows as a tetramer with the brown Pac-Man shape representing nuclease domains, the grey and black lines representing the CTD and linker, respectively, and the dark grey oval representing the GT1 domain. For simplicity each activity is illustrated by a different monomer, although whether any tetramer is active in this way is unknown. Top left, anchoring to the nucleoid through GT1 dsDNA binding activity and subsequent passage to the catalytic site. Bottom, degradation of dsRNA (left, *trans*-dsRNA; right, *cis*-dsRNA) through GT1 dsRNA binding activity. Top right, GT1-independent 5’⇾3’ exonucleolytic maturation of an mRNA where the mature terminus is protected by an RNA binding protein (RBP). **(B)** Localization of RNJ in the WTc line, favoring nucleoid localization (blue oval) anchored to thylakoid, with low dsRNA accumulation. **(C)** Same as **(B)** but for the mGT and ΔGT lines. RNJ distribution is more biased towards the stroma, and dsRNAs accumulate in granule-like bodies in the proximity of the nucleoid.

The dsRNA binding capability of the GT1 domain is most easily rationalized as a surveillance module for sense-antisense duplexes, which are expected to be common in the primary transcriptome of the chloroplast given its profusion of promoters and what appears to be mainly stochastic transcription termination. Assuming that RNase J identifies such duplexes, several questions arise. The first is how or whether it distinguishes cases where both strands are functional (e.g. *psbN* being antisense to *psbB* operon mRNAs) and spares them from degradation, and the second question is how it distinguishes the functional strand from the interfering strand, and if such a distinction exists, the third is how the targeted strand is targeted for degradation.

There are two general ways to consider such mechanistic questions. In one view, there is little mechanistic specificity, and RNase J is functioning in a “lawless” environment in competition with the splicing, editing and translational machinery. While perhaps unappealing from a regulatory point of view, the fact that chloroplast RNA is so abundant, in some cases over 80% of the cellular RNA content (63), that the many chloroplasts present in each cell are polyploid (64), and that major chloroplast proteins and RNAs generally have much longer half-lives than their bacterial counterparts (e.g. 65, 66), suggests that a highly regulated gene expression environment is neither likely nor necessary. The tolerance of easily detected dsRNA in mGT/ΔGT is consistent with such a view. A second view is that RNase J has evolved to surveil dsRNA by itself or with cellular partners in order to efficiently accomplish the tasks mentioned above. There is little evidence, however, to support this hypothesis and indeed, our own effort to seek protein partners of RNase J through a yeast two-hybrid screen was unsuccessful (unpublished results). We therefore propose that RNase J’s lack of sequence specificity is ideally suited to a generalized role in surveillance that competes sufficiently with other gene expression functions.

Regardless of the mode of surveillance, the GT1 domain is not catalytic and, in our model, needs to “feed” dsRNA to the catalytic site of RNase J. While the structure of a GT1 domain-containing RNase J has not been published, a recent study of *Helicobacter pylori* RNase J concluded that it is most active as a tetramer and that the CTD may be involved in a gating function for the catalytic site (67). We can propose that following the binding of dsRNA, the GT1 domain may subsequently position that substrate for catalysis as suggested in Fig. 13A.

In organisms where RNase J lacks the GT1 domain, this surveillance function could not occur in the same way, however there would be no impediment to the catalytic function of the enzyme based on our *in vitro* analysis (12). In this situation, RNase J would still be able to degrade asRNA, however it might be less efficient in locating them, allowing the buildup that we observe in our transgenic lines. In considering the *Chlamydomonas* RNase J that lacks a GT1 domain, it may be relevant that its chloroplast genome is arranged very differently from those of plants, with blocks of genes (“sidedness”) that might result in a much lower presence of asRNA (68). In addition, *Chlamydomonas* lacks the NEP chloroplast RNA polymerase (69) whose promoter selectivity tends to be very loose (70) and which is responsible for many seemingly unnecessary transcription start sites (1, 2). Therefore, GT1 domain-containing RNase J’s may have evolved in response to less regulated chloroplast gene expression, rather than the domain contributing to a higher specificity of RNase J.

## Supporting information

DataS1_Proteomic

DataS2_RNA-Seq

## Acknowledgments

The authors thank Drs. Yael Hacham and Guy Levin for their assistance in preparing the samples for proteomics, and the Smoler Proteomics Center at the Technion for performing the MS experiments.

## AUTHOR CONTRIBUTIONS

Kevin Baudry: Conceptualization, Data curation, Formal analysis, Investigation, Methodology, Project administration, Visualization, Writing. Sébastien Skiada: Data curation, Formal analysis, Investigation, Visualization. Carmit Burstein and Varda Liveanu: Data curation, Formal analysis, Methodology, Investigation, Visualization. Arnaud Liehrmann: Data curation. Maddie Ceminsky: Investigation. Oded Kliefeld: Data curation, Formal analysis, Investigation, Visualization. Etienne Delannoy: Formal analysis, Data curation. Alexandra Launay-Avon: Investigation. Benoît Castandet: Conceptualization, Data curation, Funding acquisition, Methodology, Project administration, Visualization, Writing. Gadi Schuster: Conceptualization, Funding acquisition, Methodology, Project administration, Visualization, Writing. David B. Stern: Conceptualization, Funding acquisition, Project administration, Writing. All authors participated in writing – review & editing.

## SUPPLEMENTARY DATA

Supplementary data are available at NAR online.

## DATA AVAILABILITY

The proteomics and RNA-Seq data underlying the analyses presented in this manuscript have been deposited with the following accession numbers:

Proteomics: [Submission in progress]

RNA-Seq: RNA-Seq and dsRIP-Seq data have been deposited in the SRA database with the project number PRJNA1345690.

Other data are provided in supplemental figures, tables, and data files. The corresponding author should be contacted regarding any data or methods not otherwise provided.

