## Supplemental_Figures-Tables for "The GT1 domain of RNase J ensures RNA quality control through dsRNA binding in Arabidopsis plastids"

A

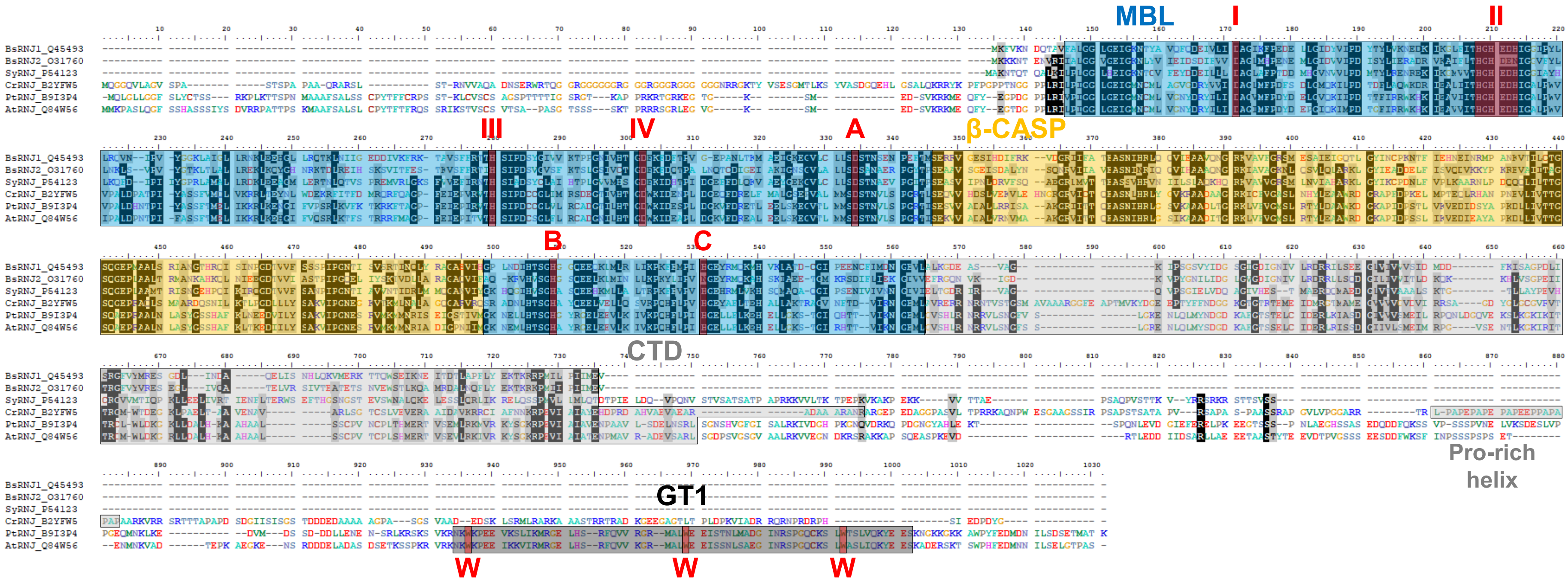


**Figure S1.** RNase J protein sequence alignment and folding comparison. **(A)** RNJ protein sequence alignment using Clustal Omega with default parameters. (BsRNJ1: Q45493; BsRNJ2: O31760; SyRNJ: P54123; CrRNJ: B2YFW5; PtRNJ: B9I3P4; AtRNJ: Q84W56). Indicated domains are MBL, Metallo-β-lactamase domain; β-CASP, β-CASP domain; CTD, C-terminal domain; GT1, GT1 domain. Conserved RNase J motifs are indicated by their respective names (I, II, III, IV, V, A, B, C). The three conserved tryptophan residues in the GT1 domain are highlighted in red and labeled “W”.

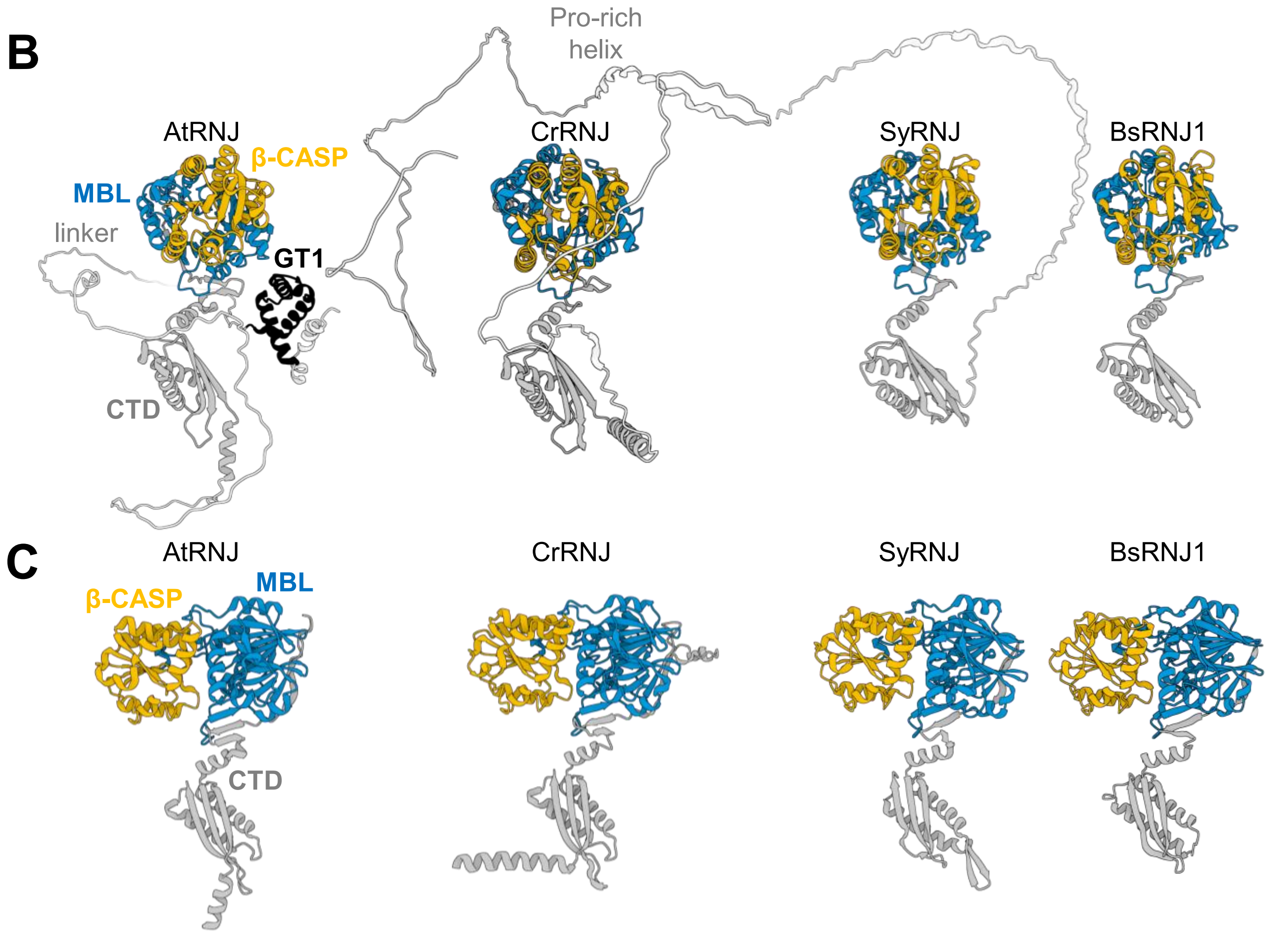


**Figure S1 (continued). (B)** AlphaFold predicted structures as accessible from the AlphaFold Protein Structure Database (https://alphafold.ebi.ac.uk/). Arabidopsis (At, AF-Q84W56-F1); Chlamydomonas (Cr, AF-B2YFW5-F1); Synechocystis (Sy, AF-P54123-F1); and Bacillus (Bs, AF-Q45493-F1). Domains are color coded as shown for Arabidopsis RNase J (AtRNJ), except for the proline-rich helix unique to *C. reinhardtii* (Cr). The N-terminal region upstream of the MBL domain is hidden for easier viewing. **(C)** Structures shown in B, rotated 90°. The N-terminal region upstream of the MBL domain and region downstream the CTD are hidden for easier viewing of the RNase J core.

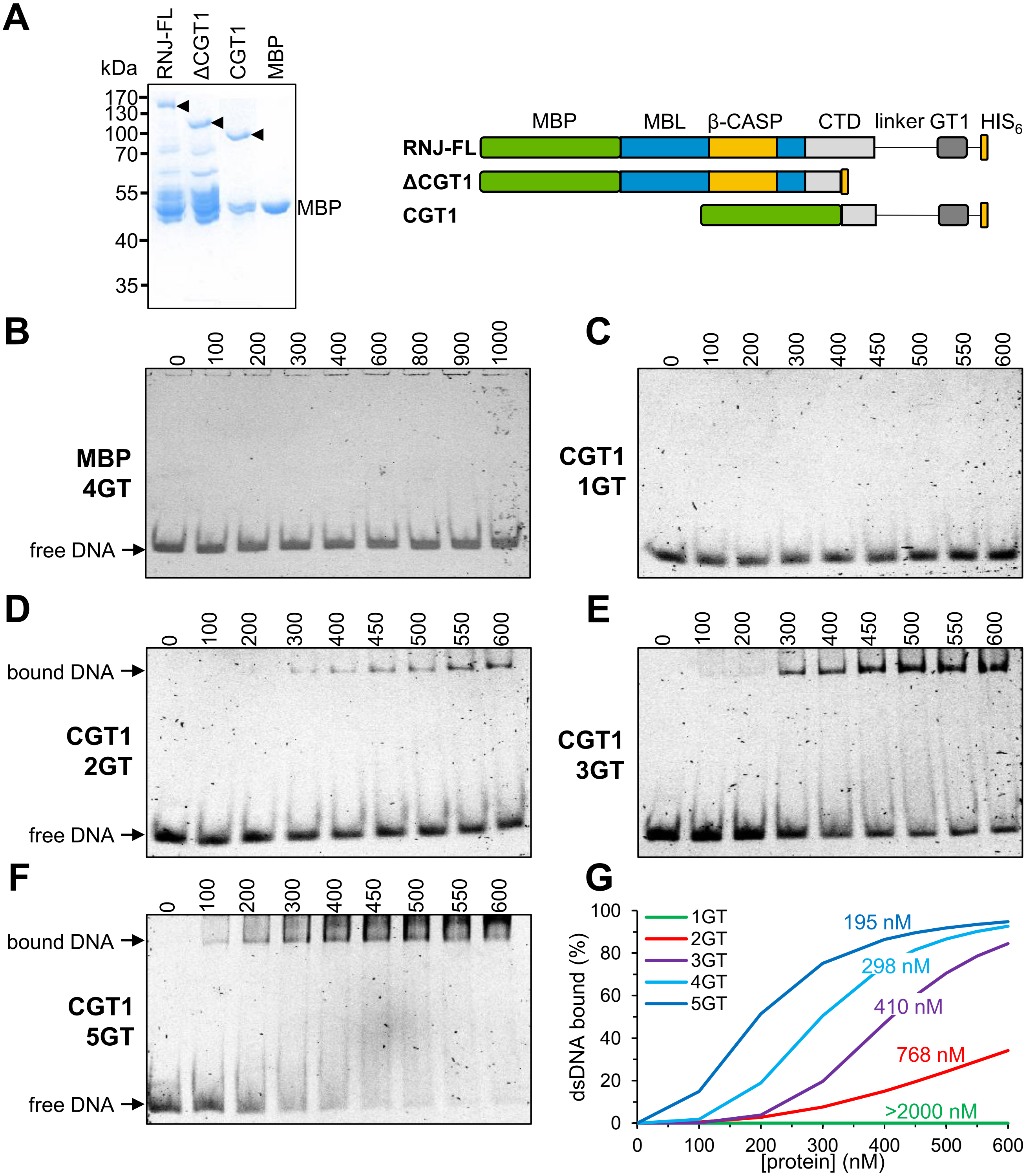


**Figure S2.** *In vitro* dsDNA binding activity of different forms of RNase J. **(A)** Expression of recombinant forms of RNase J. Left: gel analysis of recombinant proteins; right: schematic presentation of the recombinant proteins. RNJ-FL, full-length RNase J lacking the chloroplast transit peptide; ΔCGT1, truncation lacking the linker and GT1 domains; CGT1, expression of CTD and GT1 domains only. **(B)** to **(F)** and **(H)** to **(N)** Gel shift analysis. The form of recombinant protein is shown to the left of each gel, the protein concentration (nM) is indicated above each lane, and the dsDNA probe used at 100 nM is shown at the left and described in Supplemental Table 2. **(G)** Binding curves obtained by quantification of the stained gels assaying the various dsDNA probe length (corresponding to Fig 2A and panels C, D, E, and F). $K_D$ values calculated from each curve are shown.

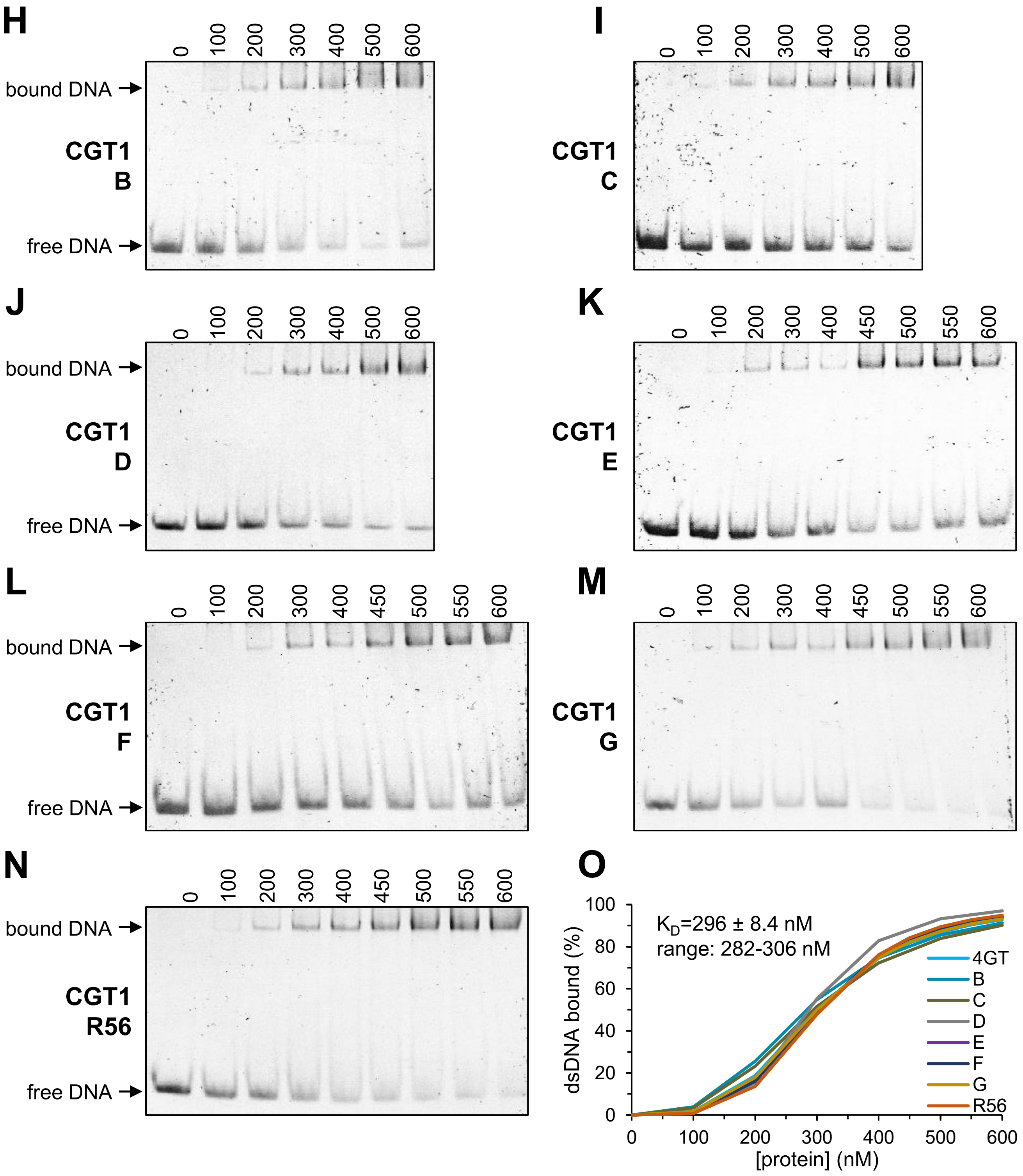


**Figure S2 (continued). (O)** As panel G, but for gels assaying dsDNA probe sequence variation (corresponding to Fig 2A and panels H to N). The average $K_D$ value is given with standard deviation.

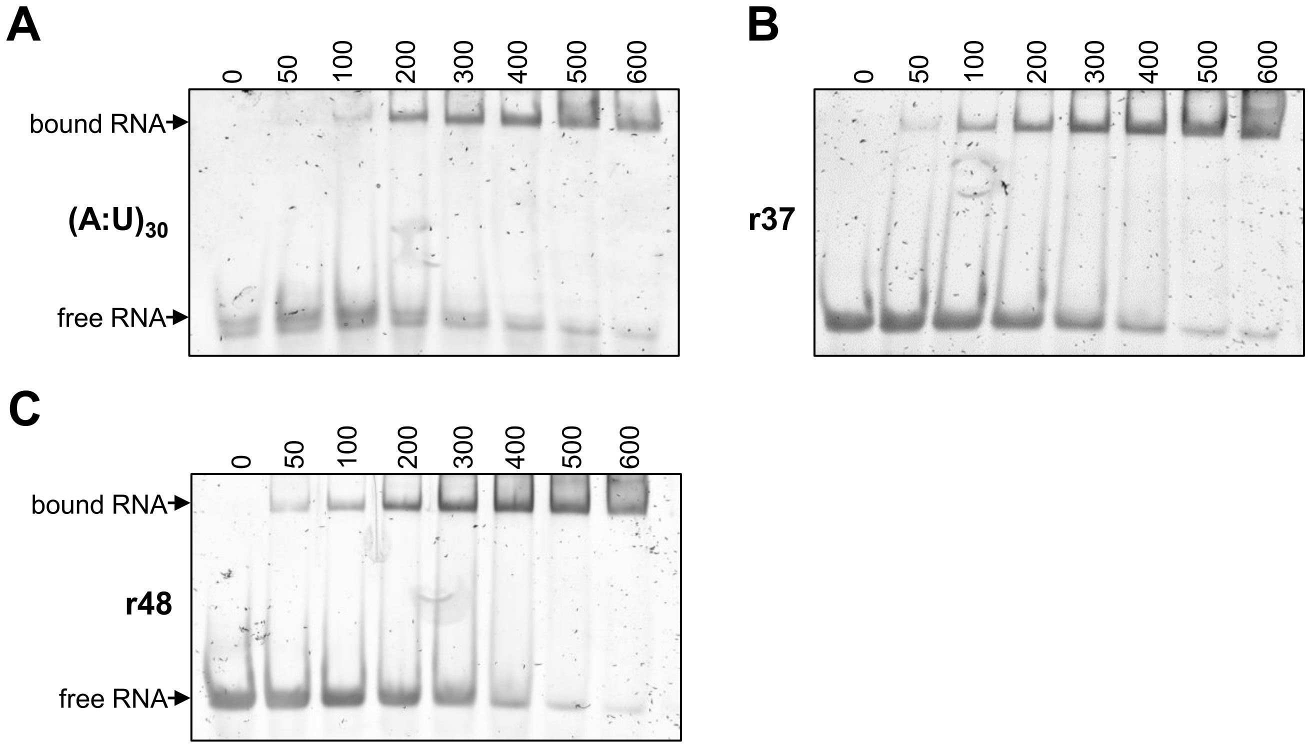


**Figure S3.** *In vitro* dsRNA binding activity. Gel shift analysis using recombinant maltose binding protein-CGT1 fusion and three different dsRNA probes at 100 nM as shown to the left of each gel and described in Supplemental Table 2. The protein concentration (nM) is indicated above each lane. Probes were **(A)** $(A:U)_{30}$, 30 bp of A and U on the two strands; **(B)** r37 and **(C)** r48, randomized dsRNAs of the given lengths.

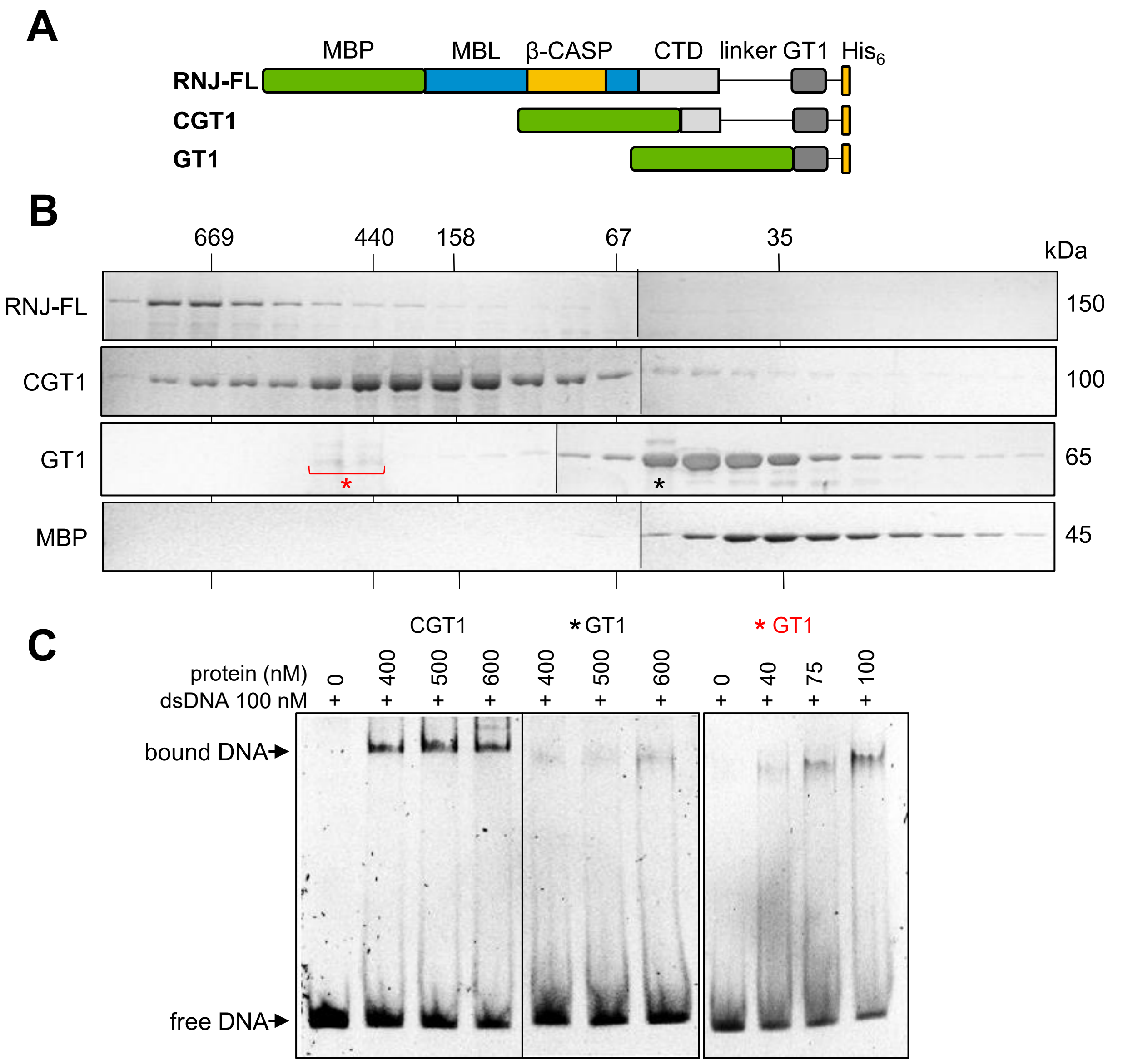


**Figure S4.** Recombinant RNJ, CGT1 but not GT1 form tetramers and this is required for high affinity binding to dsDNA. **(A)** Schematic presentation of the recombinant proteins. **(B)** Recombinant proteins purified as MBP fusions were fractionated on a Superdex 200 size exclusion column. The proteins in each fraction were detected by SDS-PAGE and Coomassie blue staining. The molecular weight of the complexes according to the elution profile of size markers is shown above, and the MW of the monomers according to SDS-PAGE is displayed to the right. The RNJ-FL and the CGT1 proteins form tetramers, while the GT1 and MBP fractionate primarily or exclusively as monomers, respectively. Similar results were obtained when the purified proteins were treated with RNase or DNase before being loaded onto the size exclusion column (data not shown). **(C)** Gel shift analysis. The form of recombinant protein and concentration are indicated above each lane; the dsDNA probe was 4GT. The results were analyzed on the same gel, and the picture is presented with the same exposure. Recombinant GT1 was from a monomeric fraction (black asterisk) or tetrameric fractions (red asterisk).

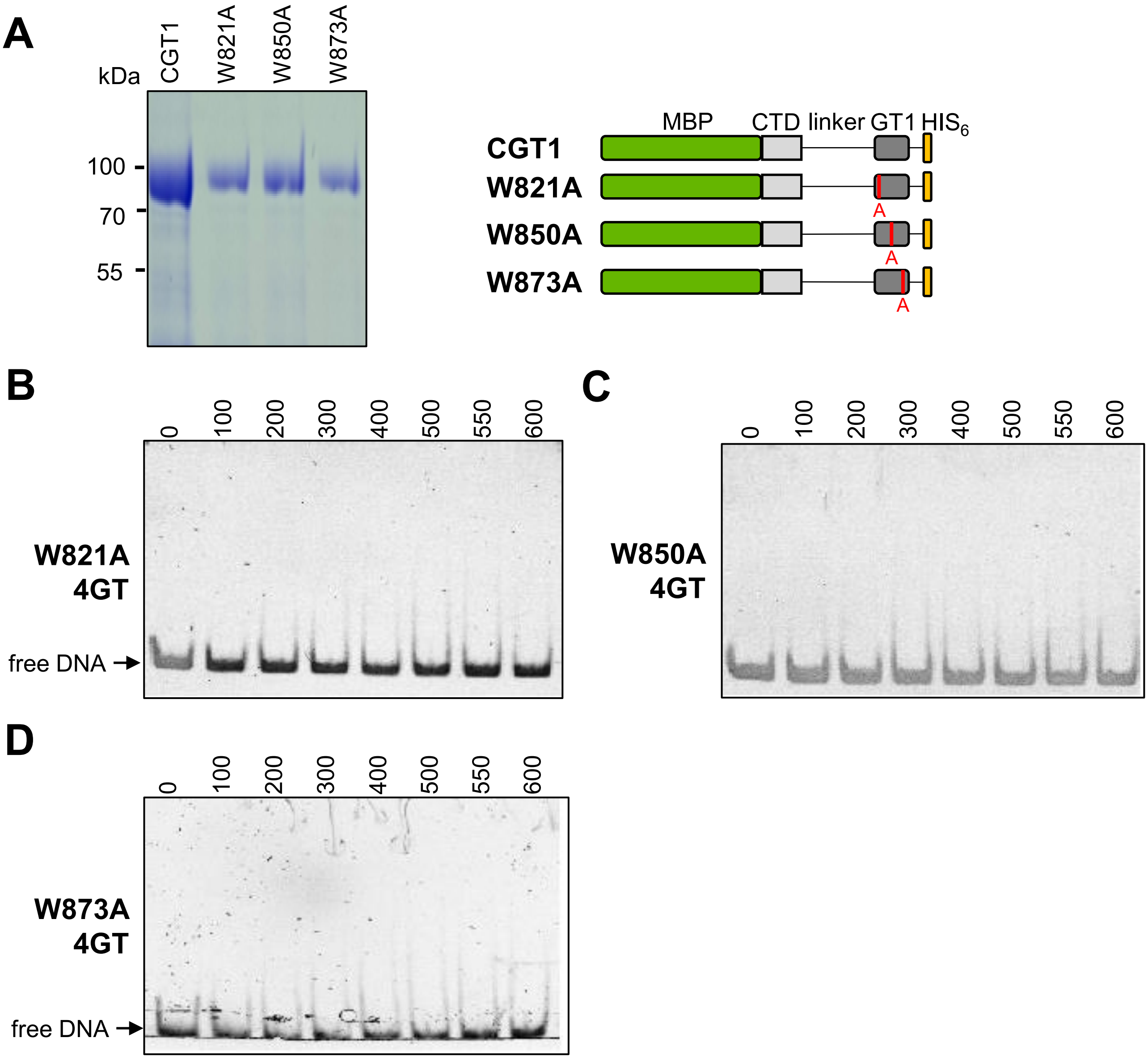


**Figure S5.** Effect on dsDNA binding of mutating Trp residues in the GT1 domain. **(A)** Expression of recombinant of wild-type and mutated CGT1 proteins. Left: gel analysis of recombinant proteins; right: schematic presentation of the recombinant proteins. **(B)** to **(D)** Gel shift assays were carried out using mutated versions of the CGT1 protein and a 4GT dsDNA probe at 100 nM. Mutations were **(B)** W821A; **(C)** W850A; and **(D)** W873A. Protein concentrations in nM are indicated above each lane.

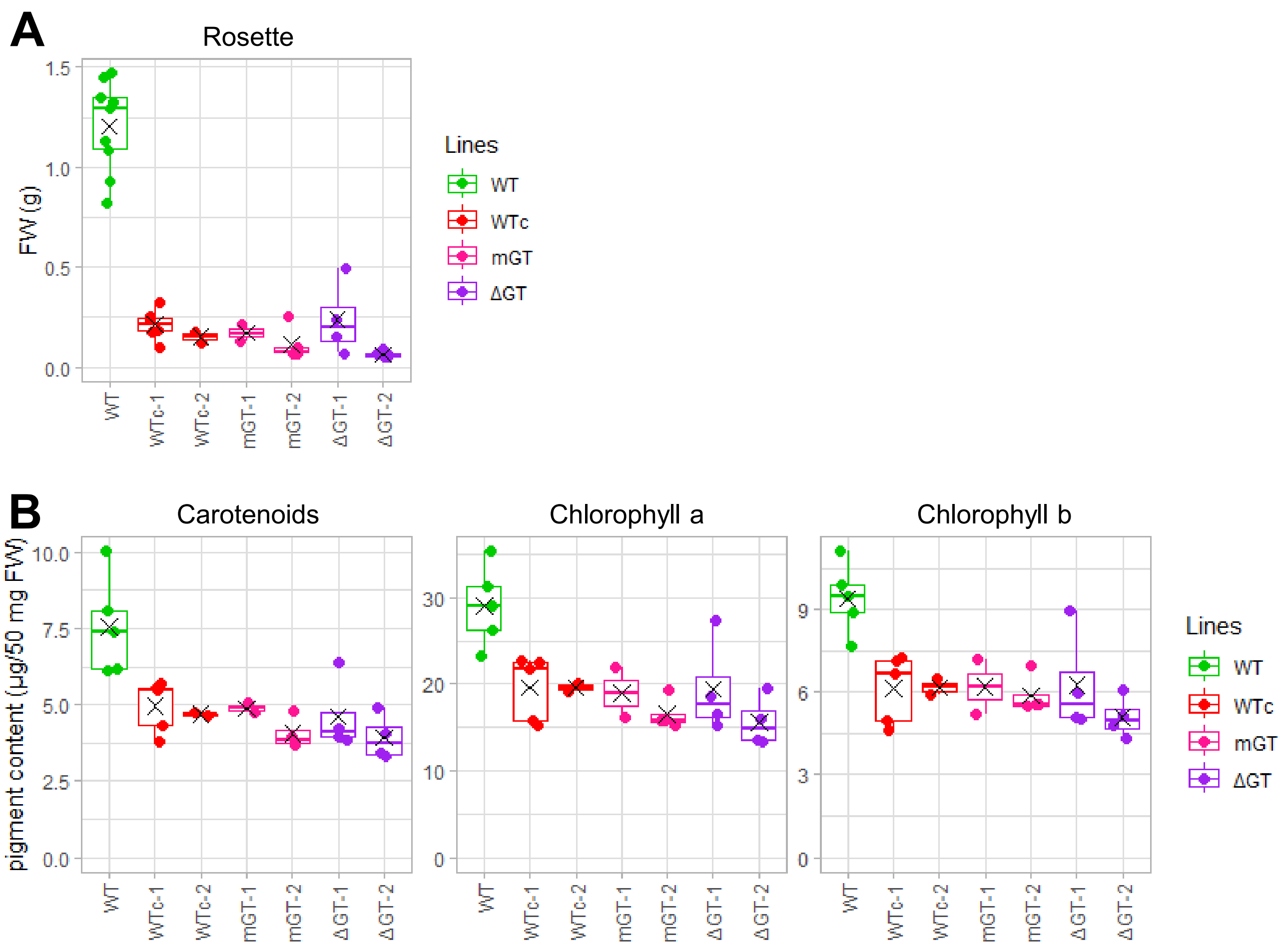


**Figure S6.** Phenotypes of complemented lines. **(A)** Average fresh weight of rosettes with roots removed, from 4-week-old plants. **(B)** Content of the pigments shown were measured in the indicated genotypes, from the same material as for Panel A.

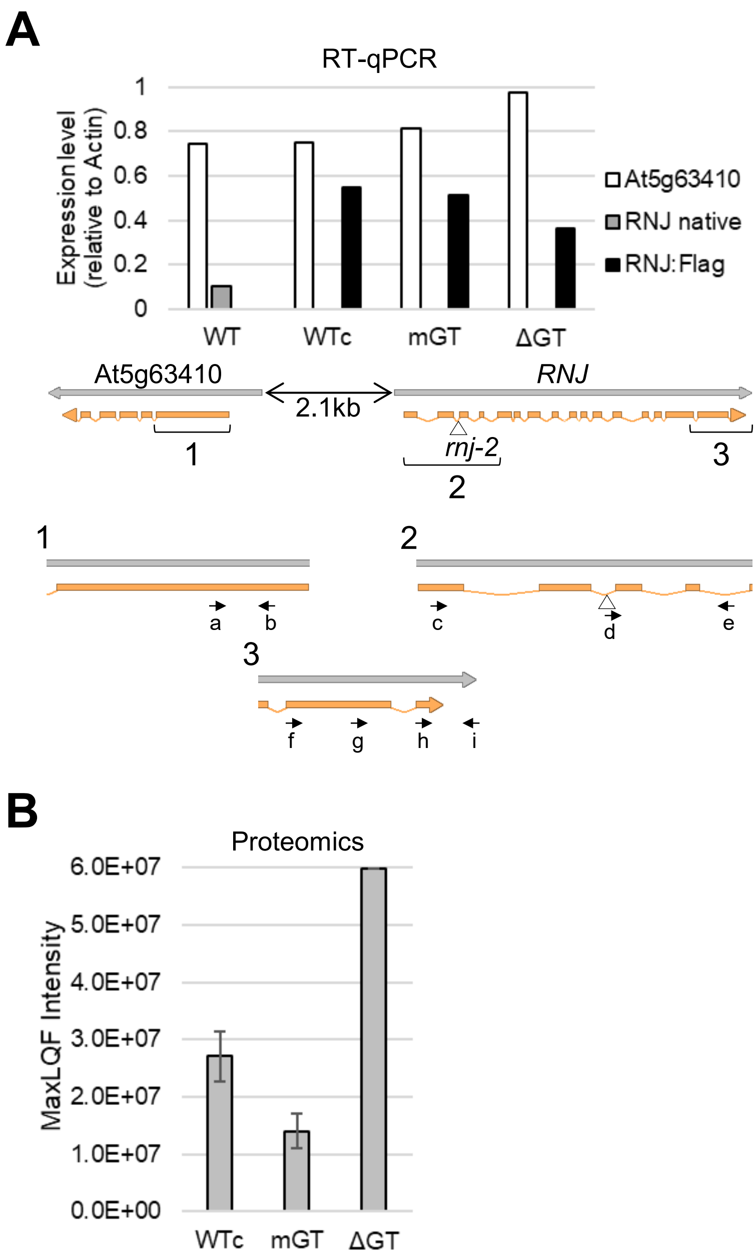


**Figure S7.** RNase J and At5g63410 expression in complemented lines. **(A)** Quantitative RT-PCR was used to query expression levels of three genes in the four genotypes indicated. At5g63410, a divergently transcribed locus upstream of *RNJ*; RNJ native, *RNJ* gene, which is disrupted at the indicated position by a T-DNA insertion in *rnj-2*; and RNJ:Flag, the respective transgenes present in the complemented lines. Regions indicated by brackets are magnified below to show primer binding sites. Region 1 shows primers (a and b) used to measure At5g63410 expression. Region 2 shows primers used to genotype the *rnj-2* T-DNA insertion, with the wild-type allele being amplified with primers c and e, and the T-DNA insertion with primers d and e. Region 3 shows primers used to measure RNJ expression, with h and i being used to measure transcripts of the native *RNJ* gene (gray bar). Transgene expression (black bars) was measured with a primer in the Flag tag and primer g (WTc, mGT) or f (ΔGT). **(B)** Normalized quantification of RNJ peptides (MaxLQF intensity) in WTc, mGT and ΔGT. Three technical replicates were obtained for WTc and mGT, and one for ΔGT.

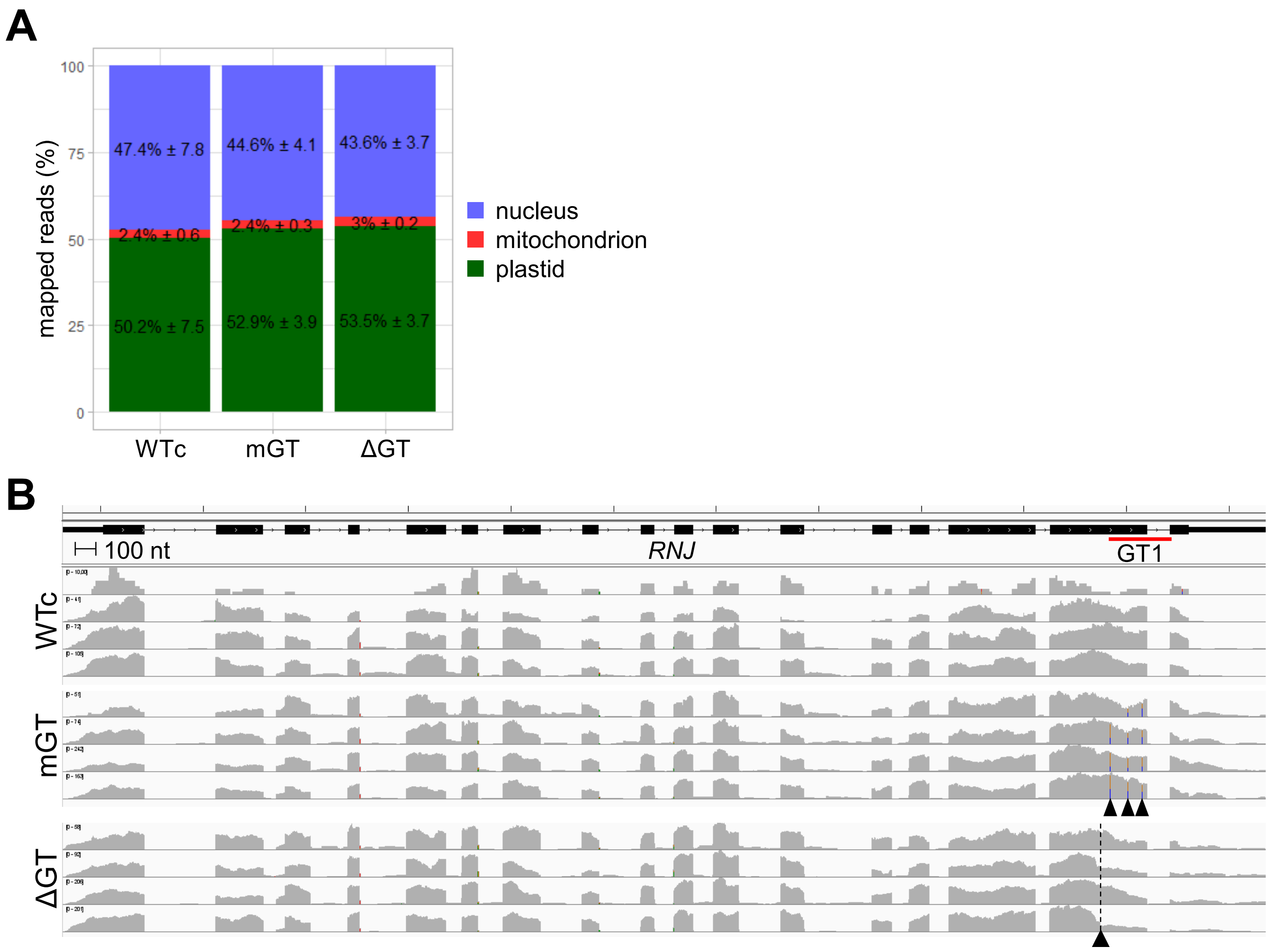


**Figure S8.** Read mapping and expression of the *RNJ* locus. **(A)** Proportion of reads from total RNA samples mapping to three cellular compartments. Reads were counted from the BAM file index using samtools idxstats, then the respective proportions were calculated. The average proportion is given with standard deviation between biological replicates. **(B)** Read coverage over the native *RNJ* locus in total RNA-Seq replicates from genotypes indicated at left, resulting in a blend of residual coverage at the T-DNA-interrupted locus and reads from the transgene. IGV was used to extract reads from four biological replicates covering chromosome 5 coordinates 25,400,317–25,406,250. The GT1 domain is indicated by the red underline beneath the gene model. The three Trp mutations in mGT line are visible by vertical dashes in the coverage indicative of mismatches and shown by arrowheads. In ΔGT, the dashed line and arrowhead indicates the truncation site of the GT1 domain.

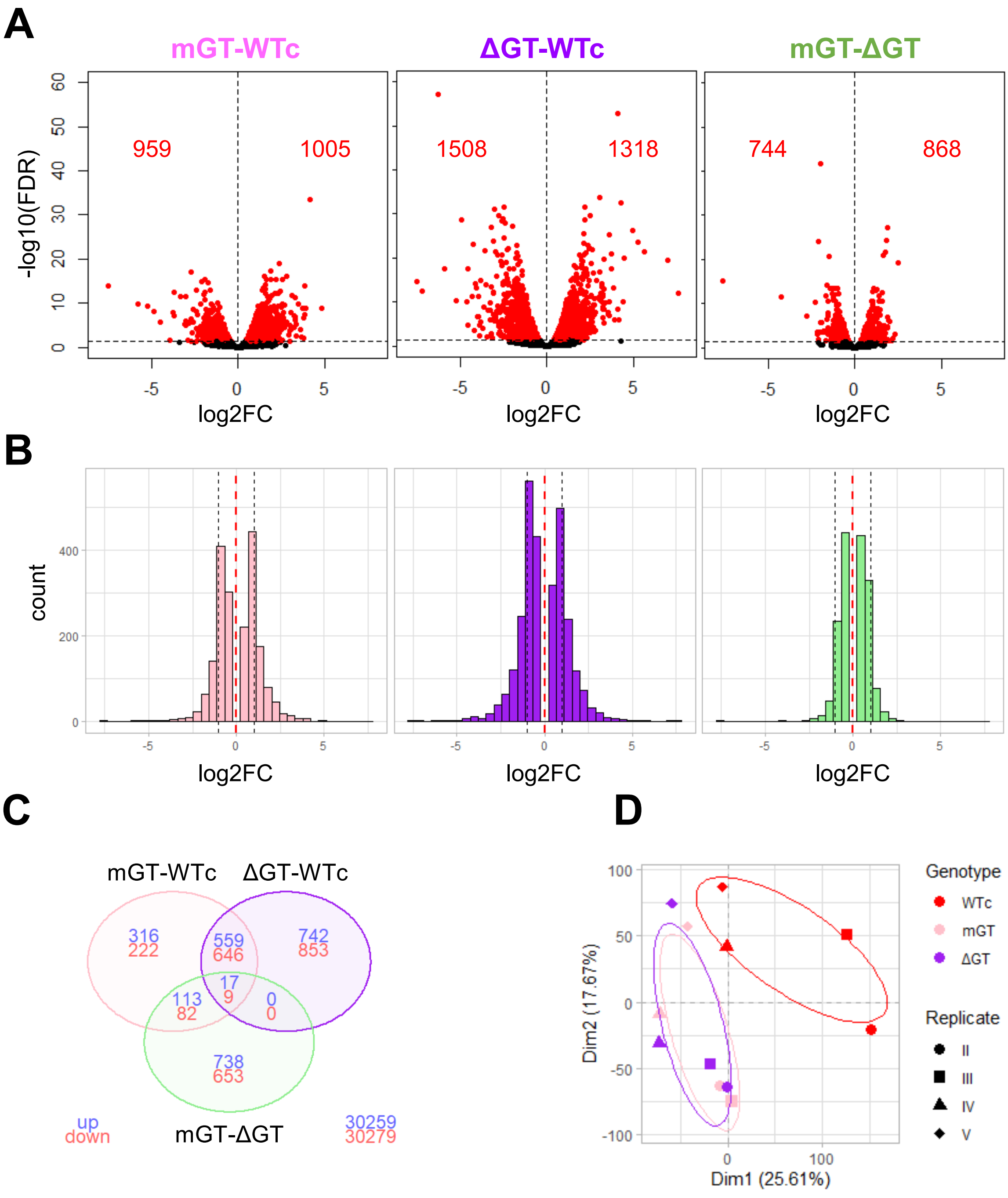


**Figure S9.** DEG identification in pairwise comparisons. **(A)** Volcano plot for DEGs identified in each pairwise comparison as indicated at the top. Each dot represents a tested gene, colored in red when difference is significant, black otherwise. Numbers on left and right side of each plot indicate quantity of significant down and up regulated DEGs respectively. **(B)** Log2 fold change distribution for identified DEGs. **(C)** Venn diagram of up (blue) and down (red) regulated DEGs. Numbers in the bottom right corner are analyzed genes that are never found as differentially expression in any comparison. **(D)** Principal component analysis (PCA) of normalized counts, colors indicate the lines and shapes are the biological replicates.

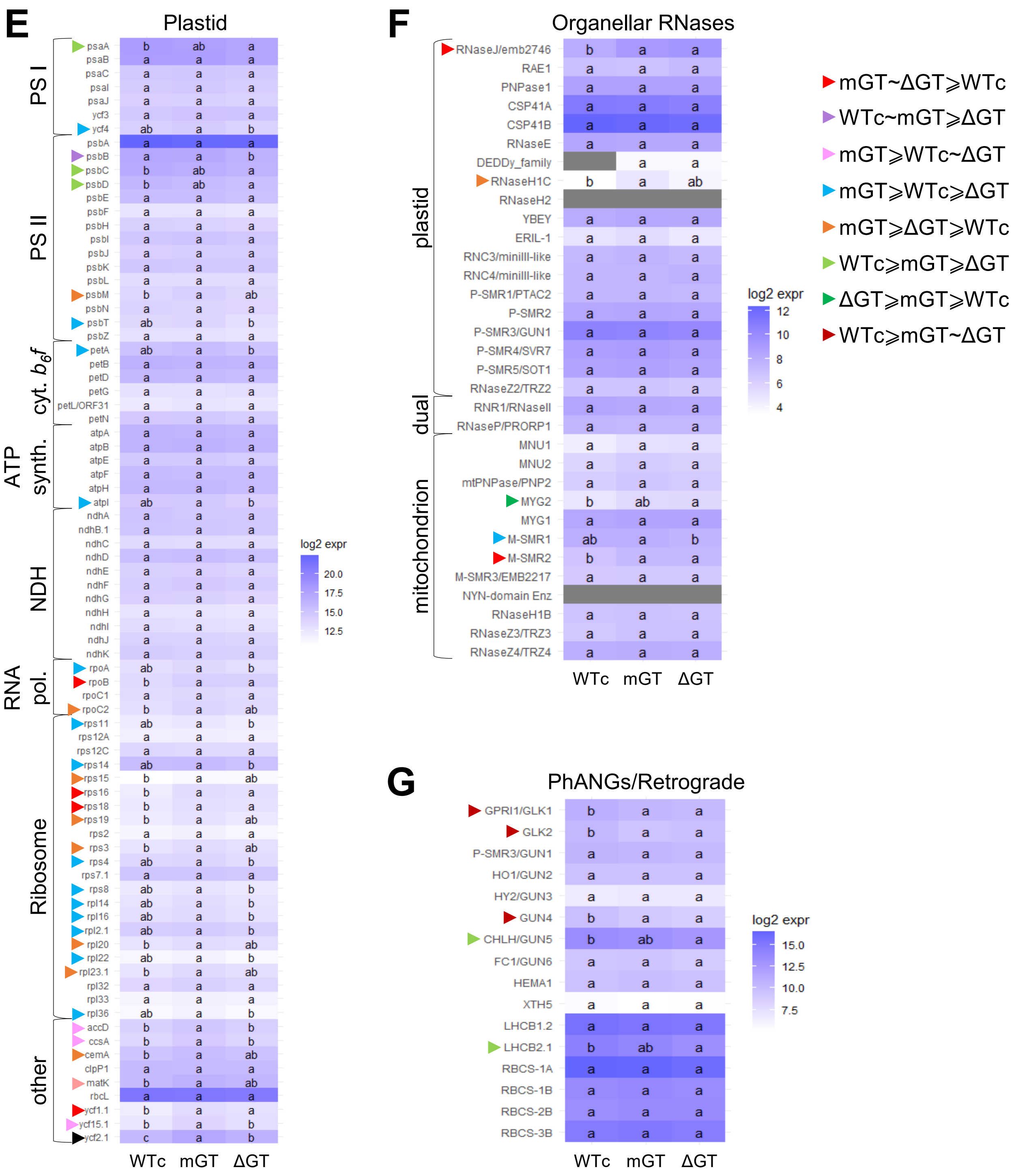


**Figure S9 (continued). (E)** Expression levels and statistical significance across genotypes for plastid protein-coding genes. The background color shading indicates the expression level obtained from RNA-Seq analysis (log2 normalized counts) in each genotype; letters indicate the significance of the difference calculated in pairwise comparisons (significance: q-value <0.05). Color-coded arrowheads signify genes with at least one significant difference, with the key of expression hierarchies shown at the top right. **(F)** Same analysis as panel E but for nucleus-encoded organellar-targeted RNases. **(G)** Same analysis as panel D, but for selected photosynthesis-associated nuclear genes (PhANGs) or genes involved in retrograde signaling.

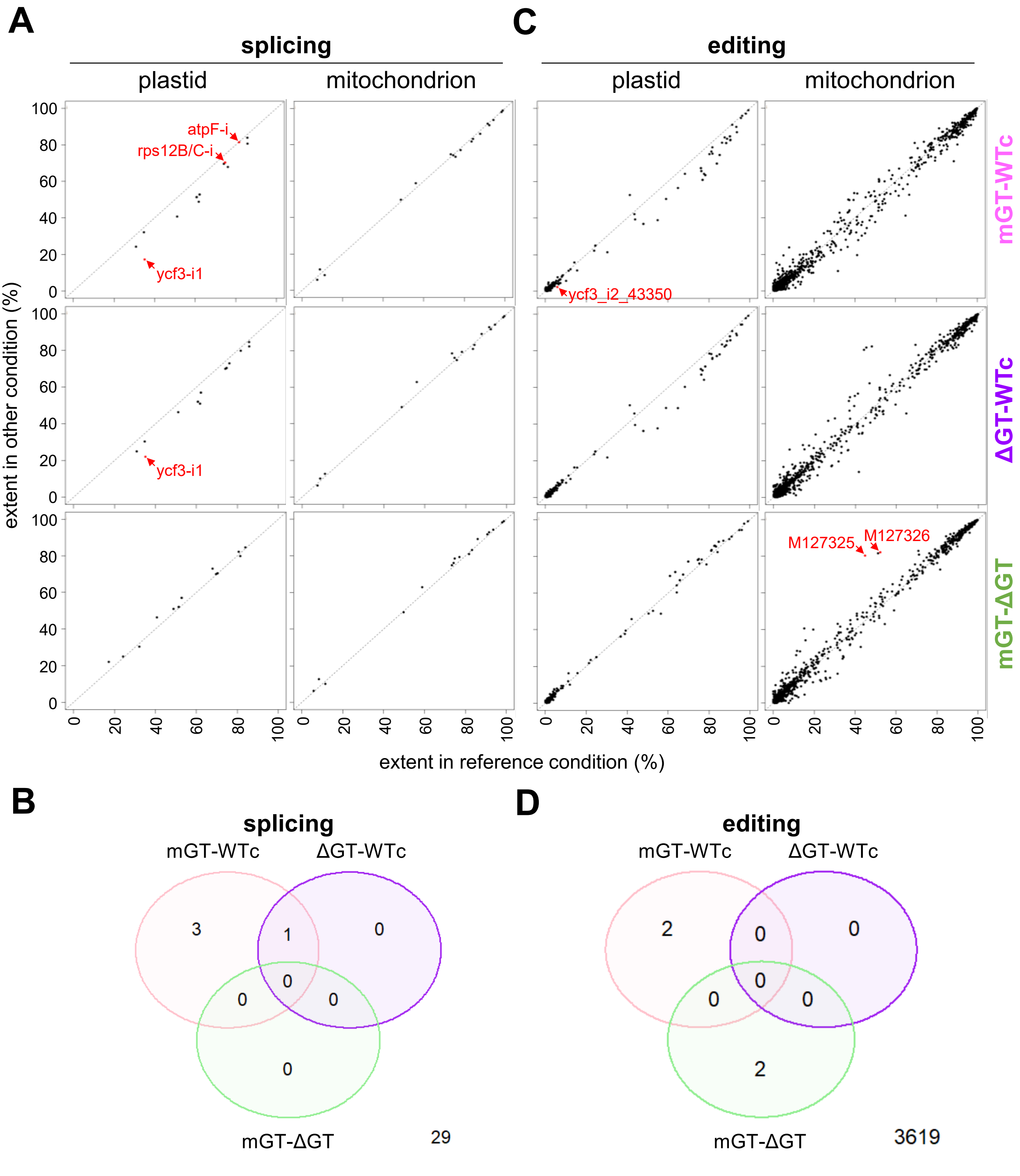


**Figure S10.** Splicing and editing of organellar transcripts. **(A)** and **(B)** results for intron splicing. **(C)** and **(D)** results for C-to-U RNA editing. **(A)** and **(C)** Each pairwise comparison is for a given organelle (top) and genotype pair (right). Dots are red when the difference is significant (red arrowheads). **(B)** and **(D)** Venn diagrams of differentially spliced introns or edited sites, respectively, corresponding to the red dots in **(A)** and **(C)**. Numbers at bottom right are total introns (29) and editing sites (3,619) not showing significant differences.

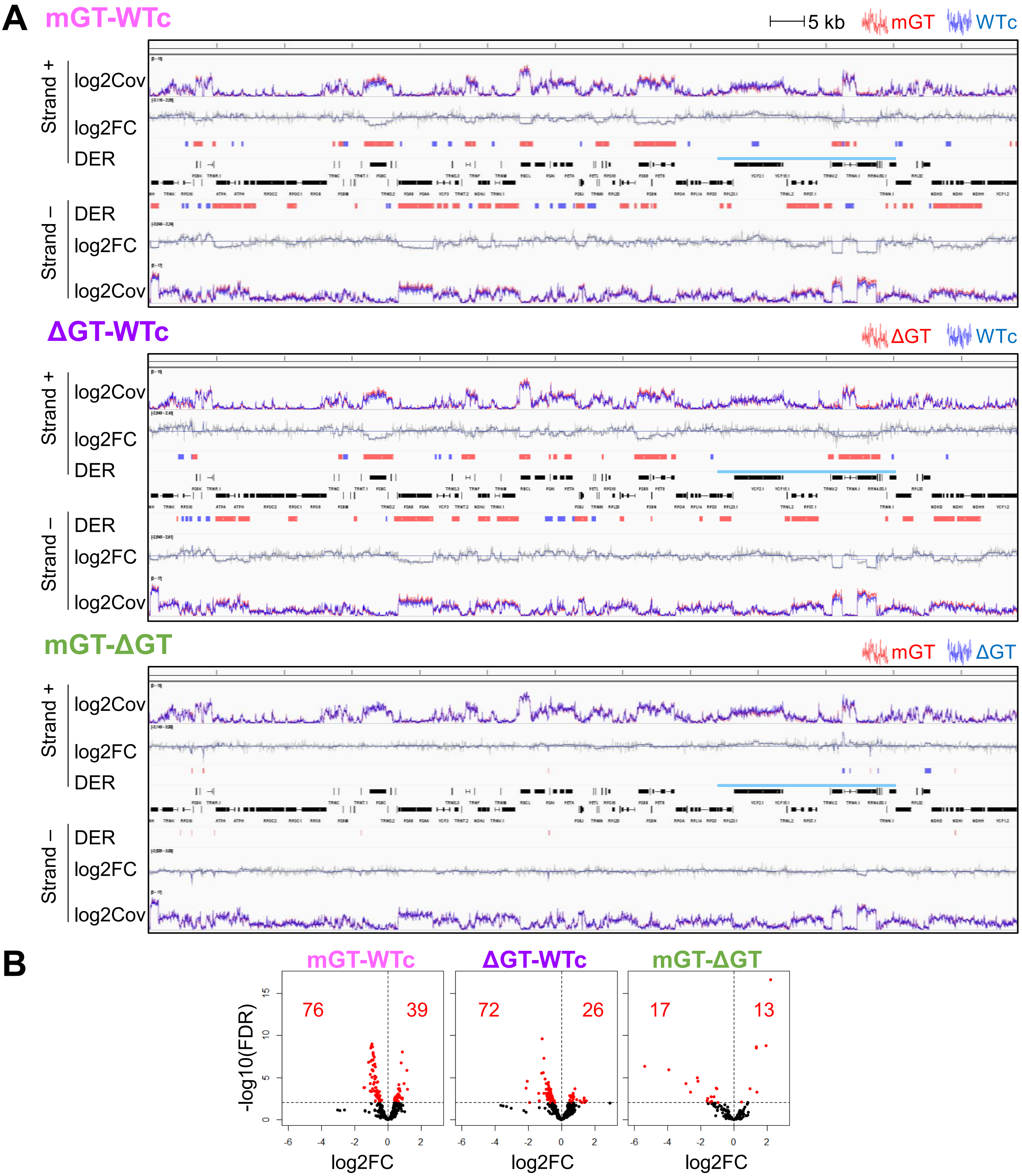


**Figure S11.** DiffSegR results and volcano plots for total RNA analysis. **(A)** Localization of DERs by strand and pairwise genotype comparison, using visualization on IGV of chloroplast genome coordinates 1-128,216, thereby excluding one copy of the inverted repeat (overlined in blue). Selected genes are annotated in black at the center. Tracks represent: log2Cov, stranded coverage on a log2 scale for each genotype, lines are color-coded according to the key on top of each panel; log2FC, per-base log2 fold change between genotypes; DER, differentially expressed regions color-coded as up-regulated (blue) or down-regulated (red). **(B)** Volcano plots for DERs identified in each pairwise comparison as described for Fig. S9A.

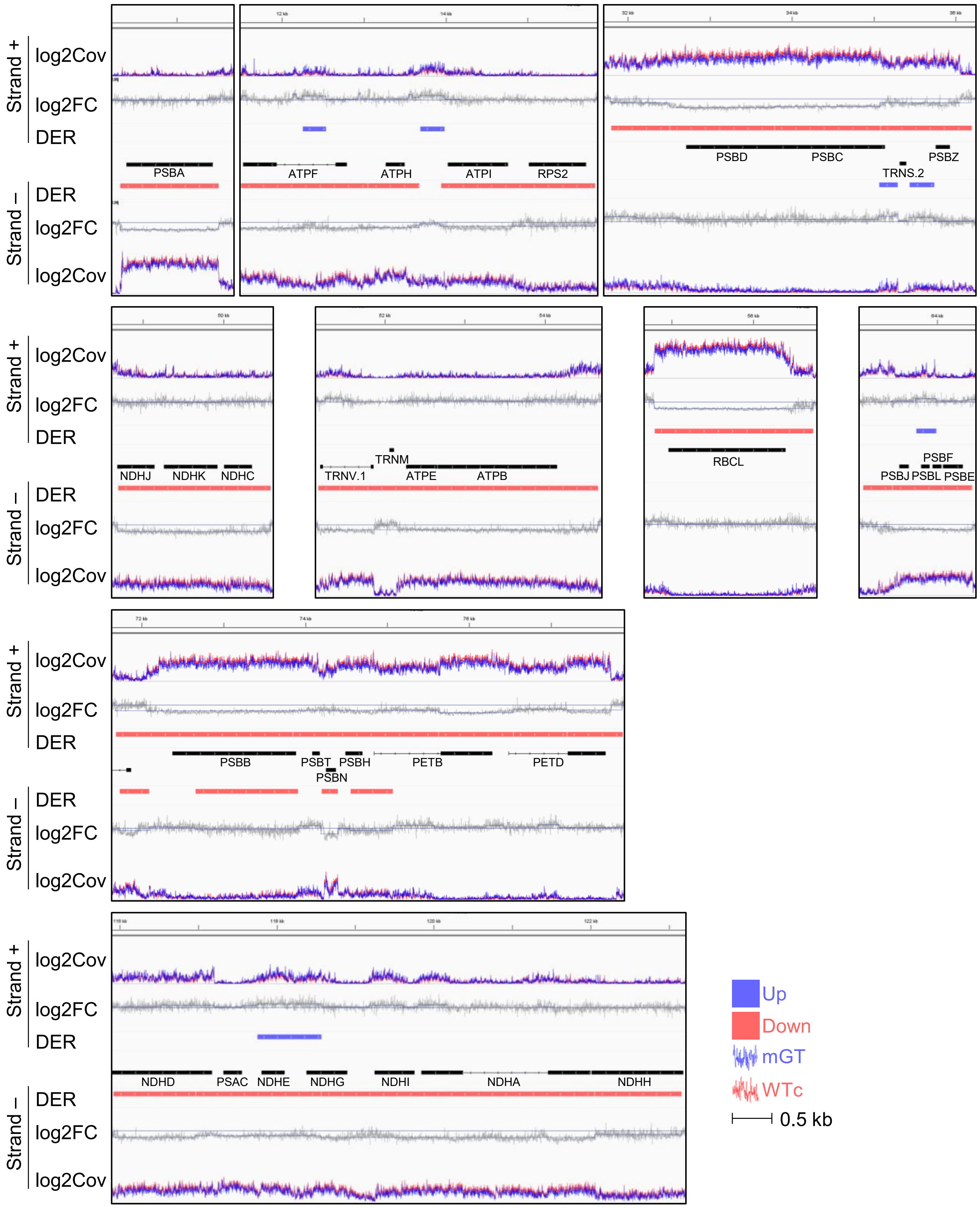


**Figure S12**. Magnification of selected DERs from the mGT-WTc comparison. Panels are as described for Fig. S11A.

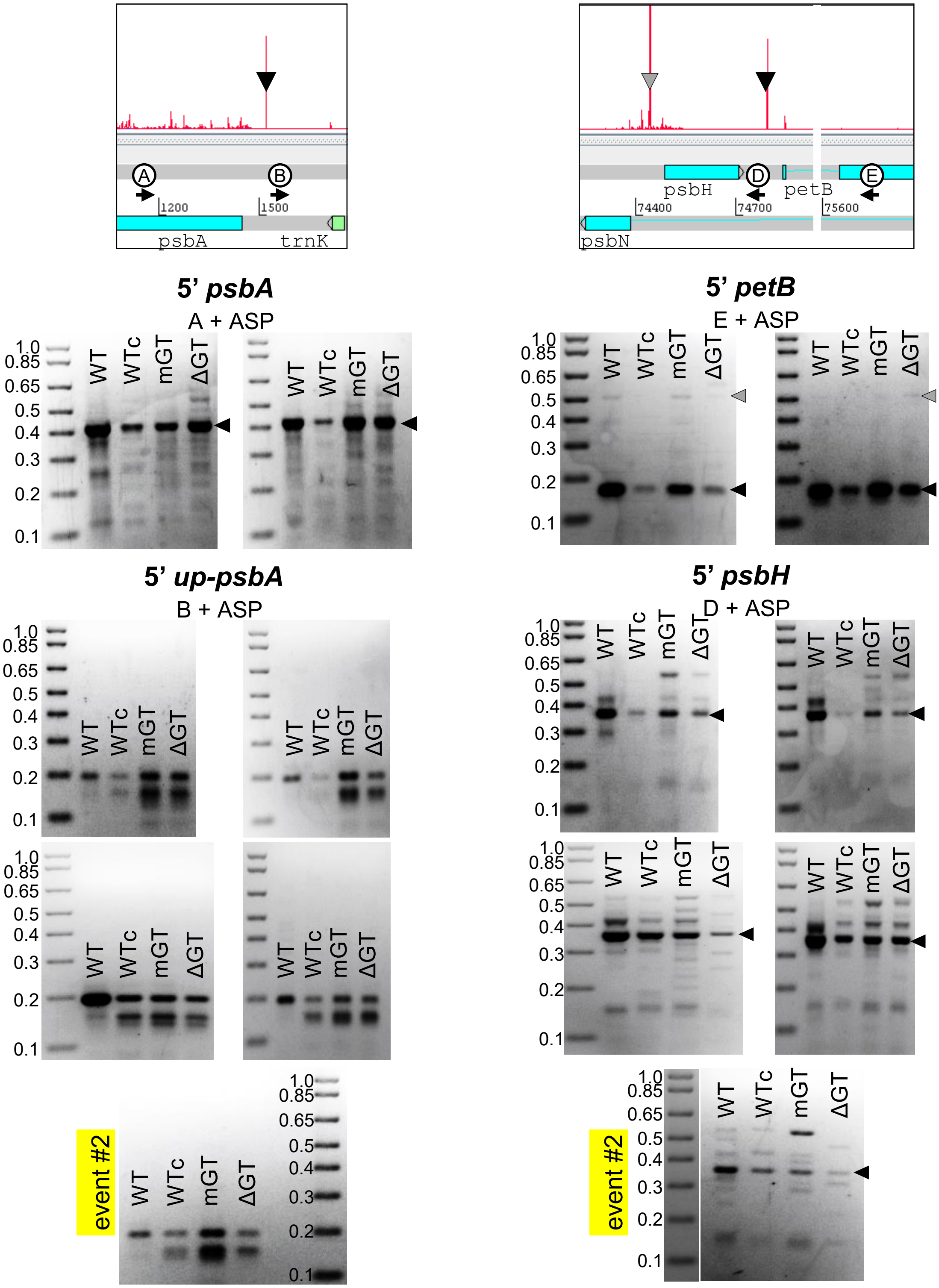


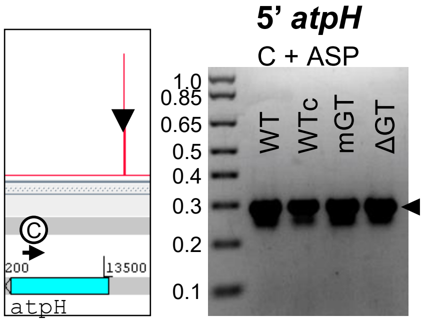


**Figure S13.** 5’ RACE analysis of *psbA*, *atpH*, *psbH* and *petB*. Primers A,B,C,D and E were used with adapter-specific primers (ASP) to amplify 5’ termini. Locations of primers are shown relative to the cognate gene. Traces indicate 5’ termini identified by Terminome-Seq (Castandet *et al.*, 2019). Arrowheads indicate the major 5’ termini, and marker sizes are in kb.

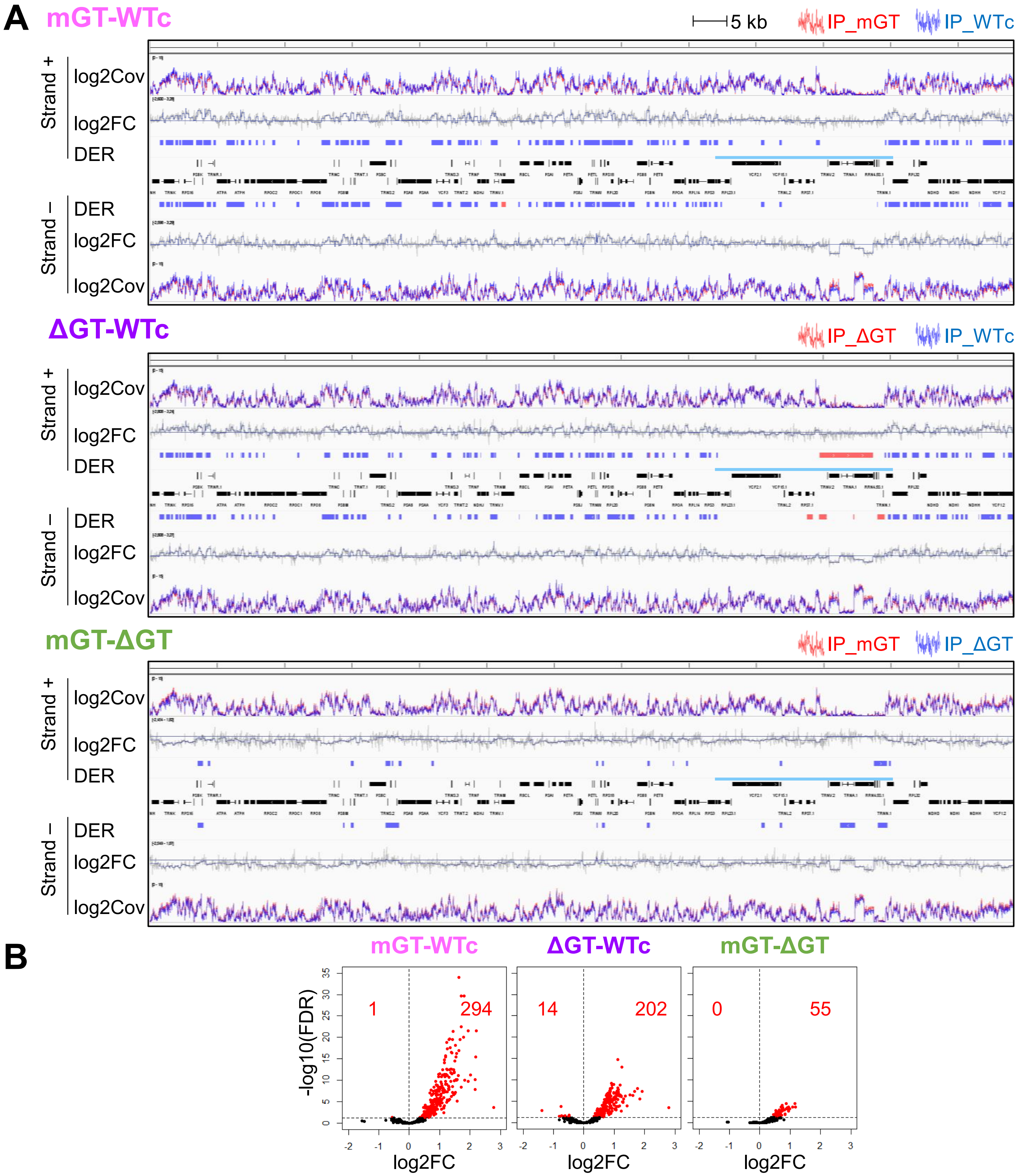


**Figure S14.** DiffSegR results and volcano plots of dsRIP-Seq data. **(A)** Localization of DERs as described for Fig. S11A. **(B)** Volcano plots for identified DERs as described for Fig. S11B.

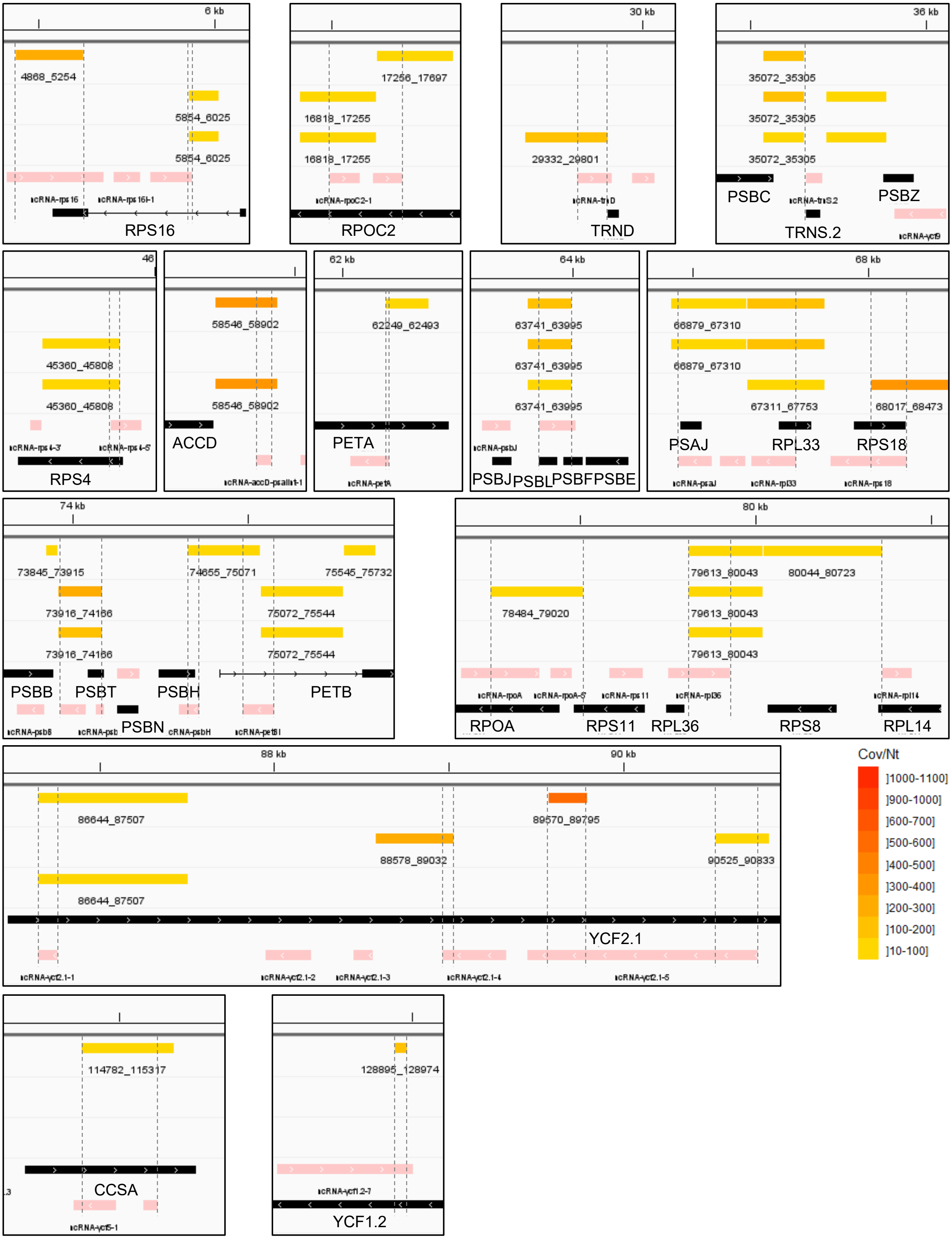


**Figure S15.** Magnification of *trans*-dsRNAs overlapping with known ncRNAs. Magnification of boxes in Fig. 11A, showing 34 intersections between 26 trans-dsRNA and 30 ncRNA. Dashed lines indicate overlapping regions.

**Table S1.** Primers used in this article

| Name | Sequence | Comments |
|---|---|---|
| **Genotyping** | | |
| rnj2_LP | CCCTCTCGCTTTGTCCCTAT | rnj-2 genotyping |
| rnj2_RP | CGCAAGCTGAGGTACATACA | rnj-2 genotyping |
| LB1 | GCCTTTTCAGAAATGGATAAATAGCCTTGCTTCC | rnj-2 genotyping |
| AtRNJ_ex16_F | ACTGCGGCATCAACATACAC | transgene genotyping |
| FLAGtag_R | TGTCATCGTCATCCTTGTAGTC | transgene genotyping |
| **RT-qPCR** | | |
| AtActin1_F | TCTTGATCTTGCTGGTCGTG | ACT1 (At2g37620) gene expression |
| AtActin1_R | GAGCTGGTTTTGGCTGTCTC | ACT1 (At2g37620) gene expression |
| At5g63410_qpcr_F | AGCAACACCATCGATGAACA | At5g63410 gene expression |
| At5g63410_qpcr_R | GACGACCCCAAATCCCTAGA | At5g63410 gene expression |
| AtRNJ_ex17_qPCR_F | GAGAAGCAAGACGAGTTGGC | native RNJ (At5g63420) gene expression |
| AtRNJ_3UTR_qPCR_R | GGGATTTGGATTCTGCGTTGA | native RNJ (At5g63420) gene expression |
| AtRNJ_GT1_qPCR_F | GAGGAGAGCTGCACAGTAGA | WTc, mGT transgene expression |
| FLAGtag_qPCR_R | AGCCTTGTCATCGTCATCCT | WTc, mGT transgene expression |
| AtRNJ_ex16_F | ACTGCGGCATCAACATACAC | ΔGT transgene expression |
| FLAGtag_R | TGTCATCGTCATCCTTGTAGTC | ΔGT transgene expression |
| **5'RACE** | | |
| 5'RACEAdaptor | AUAUGCGCGAAUUCCUGUAGAACGAACACUAGAAGAAA | RNA adaptor ligated |
| ASP | AATTCCTGTAGAACGAACACTAG | Adaptor Specific Primer |
| 5'RACE_AtPsbA | AGCCATTCATCAACGGATGC | A, amplify psbA main 5' (match psbA CDS) |
| 5'RACE_AtPsbA.up | GCCATGTCAACCAATGTAAAATG | B, amplify psbA secondary 5' (match upstream psbA main 5') |
| 5'RACE_AtAtpH | GCGCTAATGCTACAACCAGGCC | C, amplify atpH 5' (match atpH CDS) |
| 5'RACE_AtPetB.exon | ACATGCGGAGGAACATATTT | D, amplify petB main 5' (match petB exon2) |
| 5'RACE_AtPetB.up | GGCTAAAATTGTTAGACCTGCAT | E, amplify psbH main 5' (match upstream petB main 5') |
| **Cloning of the RNJ cDNA to a bacterial expression plasmid** (restriction sites are underlined) | | |
| RNJ_R1 | GCCTCGAGTTAAGACGCAGGTGTGCCTAG | 3' primer to amplify the RNJ, CGT1 and GT1 cDNA to clone into the expression vector |
| RNJ_F1 | GCGCGAATTCGATGATGAAACCTGCTTCTTTGCAAGGT | 5' primer to clone the RNJ without the chloroplast transit peptide. |
| CGT1_F1 | GCGCGAATTCGTGCATGTGGCTTGACAAAGGAAGACTTTTA | 5' primer to clone the CGT1 cDNA to the bacterial expression vector |
| GT1_F1 | GCGCGAATTCGAGGAAGAACAAATGGAAACCGGAGGAG | 5' primer to clone the GT1 cDNA to the bacterial expression vector |
| ΔCGT1_F | AAAGGAAAAATAAGAATCACAACGCGGCGAAGCCCGGGACAATGCAAATCTCTC | 5' primer to site-delete the CGT1 domain |
| ΔCGT1_R | GAGAGATTTGCATTGTCCCGGGCTTCGCCGCGTTGTGATTCTTATTTTTCCTTT | 3' primer to site-delete the CGT1 domain |
| **Site-directed mutagenesis of the conserved tryptophans in the bacterial expressed CGT1** | | |
| W1-F | GAAGAACAAAgccAAACCGGAGGAG | |
| W1-R | CTCACACGTTTTGGTGATG | |
| W2-F | AATGGCATTGgccGAAGAGATCTCTTCAAATC | |
| W2-R | CTACCTTTCACCACTTGAAATC | |
| W3-F | CAAATCTCTCgccGCATCACTTATTCAGAAATACGAGGAG | |
| W3-R | CATTGTCCCGGGCTTCGA | |

**Table S2.** DNA and RNA probes used in EMSA experiments

| Name | Sequence | length | nucleic |
|---|---|---|---|
| 1GT | 1x(GTGTGGTTAATATG) | 14 | DNA |
| 2GT | 2x(GTGTGGTTAATATG) | 28 | DNA |
| 3GT | 3x(GTGTGGTTAATATG) | 42 | DNA |
| 4GT | 4x(GTGTGGTTAATATG) | 56 | DNA |
| 5GT | 5x(GTGTGGTTAATATG) | 70 | DNA |
| B | 4x(GTGTGAGTGTTATG) | 56 | DNA |
| C | 4x(GTGTGGTTCCTATG) | 56 | DNA |
| D | 4x(GAGAGAGAGAGAGA) | 56 | DNA |
| E | 4x(TATAGGTTAAGGTG) | 56 | DNA |
| F | 4x(GTGTGGGTGTTATG) | 56 | DNA |
| G | 4x(CTCTGGTTCCCTCT) | 56 | DNA |
| R56 | AATACCAACCACCCTGTTAAATGCCATCGGACTCAGTAATCTATGATATGCTCTCT | 56 | DNA |
| A30 | 30x(A) | 30 | RNA |
| U30 | 30x(U) | 30 | RNA |
| N56 | GAUGUGGUUAAUAUGGUGUGGUUAAUAUGGUGUGGUUAAUAUGGUGUGGUUAAUAUG | 56 | RNA |
| r37 | AAUUUCUAAUAAAUACAUUUUUCCUCUCACACACUUU | 37 | RNA |
| r48 | AUUGUAUCCUUAACCAUUUUUUUUUGACACGAGGAACUCAUCAUGCUC | 48 | RNA |

**Table S3.** Selected enrichment analysis results
Lists of overrepresented terms identified for each pairwise comparison as indicated at the top and by annotation system (GO, top; KEGG, middle; MapMan, bottom). For each annotation, three DEG lists were analyzed independently: Up-regulated (log2FC >0, q-value <0.05) on top; Down-regulated (log2FC <0, q-value <0.05) in the middle; all identified DEGs (q-value <0.05) at the bottom. For the KEGG annotation, pathway term are in bold, KO terms are in plain type. Words written in blue or red denote terms shared by both GT1-deficient comparisons with WTc across annotations for up and down-regulated gene lists, respectively.

| Annotation | List | mGT-WTc | ΔGT-WTc | mGT-ΔGT |
|---|---|---|---|---|
| GO | Up | auxin (import, homeostasis)<br>cell wall biog., cutin biog., cell wall loosening, xyloglucan metab.<br>unidimensional cell growth<br>neg. reg. DNA duplication<br>long chain fatty acid metab.<br>zygote asymmetric cytokinesis in embryo sac<br>resp. to (brassinosteroid) | auxin import<br>cell wall biog., cell wall modif., xyloglucan metab., pectin catab.<br>brassinosteroid mediated sign.<br>undimensional cell growth, root hair cell dvp.<br>long chain fatty acid metab.<br>nitrate transp., cell resp. to nitric oxide<br>resp. to (oomycetes, fungi, SA, brassinosteroid) | flavonoid biosynth., anthocyanin containing compound metb.<br>starch biosynth, glycogen biosynth., glucosinolate biosynth<br>induced systemic resistance<br>stomatal complex morphogenesis<br>resp. to (bacterium, insect, JA, water deprivation, wounding, hypoxia) |
| GO | Down | circadian rhythm<br>leaf senescence<br>pos. reg. DNA transcription<br>sporopollenin biosynth.<br>cell resp. to iron ion starvation<br>resp. to (ABA, JA, salt, cold, wounding, water deprivation, hypoxia) | circadian rhythm<br>pos./neg. reg. DNA transcription<br>photomorphogenesis, de-etiolation, chll catab., red/far red light sign.<br>flavonoid biosynth., anthocyanin containing compound metb.<br>hyperosmotic salinity resp.<br>resp. to (ABA, JA, salt, cold, water deprivation, wounding, desiccation, high light, UV-B, hypoxia) | carbohydrate transmb transp. activity |
| GO | All | cell wall biog., phloem/xylem histogenesis, cell wall loosening, xyloglucan metab.<br>circadian rythm<br>pos. reg. DNA transcription<br>SCF-dep proteasomal ubiquitin-dep protein catab.<br>resp. to (ABA, auxin, JA, cold, salt, wounding, water deprivation) | ABA sign.<br>cell wall biog., xyloglucan metab., cell wall loosening, pectin catab.<br>circadian rhythm, entrainment of cicadian clock<br>red/far red light sign.<br>sterol biosynth<br>defense reponse<br>resp. to (ABA, auxin, JA, SA, fungi, oomycetes, bacterium, cold, salt, light stimulus, water deprivation, wounding) | flavonoid biosynth., anthocyanin containing compound metb.<br>glycogen biosynth., glucosinolate biosynth<br>cinnamic acid biosynth.<br>resp. to (bacterium, insect, JA, oxidative stress, water deprivation, wounding, hypoxia) |
| KEGG (pathway) | Up | **DNA replication**<br>**fatty acid elongation**<br>**pentose and glucuronate interconversions**<br>tubulin alpha<br>ABA recept. PYR/PYL<br>3-ketoacyl-coA synth.<br>xyloglucan:xyloglucosyl transferase, pectate lyase | **fatty acid elongation**<br>**plant-pathogen interaction**<br>**plant hormone sign.**<br>**phenyl propanoid biosynth.**<br>tubulin beta<br>ABA recept. PYR/PYL<br>3-ketoacyl-coA synth.<br>xyloglucan:xyloglucosyl transferase<br>peroxidase<br>SAUR family | **2nd metabolites biosynth.**<br>**nulceotide sugar biosynth.**<br>**flabonoid biosynth.**<br>**galactose metab., glucosinolate biosynth.**<br>**starch and sucrose metab.**<br>LRR recept. S/T kinase ERECTA<br>glucose-1-P adenyltransferase |
| KEGG (pathway) | Down | **circadian rhythm**<br>EREB-like factor<br>protein phosphatase 2C (PP2C) | **circadian rhythm**<br>**2nd metabolites biosynth.**<br>**carotenoid biosynth., flabonoid biosynth.**<br>**starch and sucrose metab.**<br>EREB-like factor<br>protein phosphatase 2C (PP2C) | **MAPK sign.**<br>SAUR family<br>MFS transporter, SP family, ERD6-like sugar transporter |
| KEGG (pathway) | All | **plant hormone sign.**<br>ABA recept. PYR/PYL<br>3-ketoacyl-coA synth.<br>EREB-like factor<br>protein phosphatase 2C (PP2C)<br>xyloglucan:xyloglucosyl transferase | **circadian rhythm**<br>**2nd metabolites biosynth.**<br>**plant-pathogen interaction**<br>**plant hormone sign.**<br>**MAPK sign.**<br>**phenyl propanoid biosynth.**<br>**starch and sucrose metab.**<br>3-ketoacyl-coA synth.<br>EREB-like factor<br>protein phosphatase 2C (PP2C)<br>xyloglucan:xyloglucosyl transferase<br>peroxidase | **2nd metabolites biosynth.**<br>**flabonoid biosynth.**<br>**galactose metab., glucosinolate biosynth.**<br>**MAPK sign.**<br>**pentose and glucuronate interconversions**<br>**starch and sucrose metab.**<br>phenylalanine ammonia lyase<br>xyloglucan:xyloglucosyl transferase |
| MapMan | Up | cell division (DNA replication, cell cycle metaphase to anaphase)<br>cell wall org. (pectin.pectate lyase, cell wall prot.hydroxyproline glycoprot. activities)<br>cytoskeleton (alpha-beta-tubulin heterodimer)<br>ER fatty acid elongase. 3-ketoacyl-CoA synth.<br>glycosyltransferase<br>ABA perception, sign.PYL/RCAR<br>TKL kinase superfamily (LRR-III) | cell wall org. (pectin, cell wall prot.hydroxyproline glycoprot. activities)<br>cytoskeleton (alpha-beta-tubulin heterodimer)<br>ER fatty acid elongase. 3-ketoacyl-CoA synth.<br>glycosyltransferase<br>ABA perception, sign.PYL/RCAR<br>TKL kinase superfamily<br>solute transp. (MFS superfamily (NRT1/PTR)) | cell wall org. (hemicellulose.heteromannan)<br>starch metab. (ADP-glucose pyroP complex)<br>cytoskeleton (kinesin)<br>phenolics (flavonoid biosynth.)<br>glucosinolate biosynth<br>TKL kinase superfamily (LRR-XIII) |
| MapMan | Down | cell wall org. (sporopollenin)<br>chll catab.<br>calcium dep. sign. (CBL-CIPK)<br>circadian clock (evening element reg)<br>ABA (conjugation, perception and sign.)<br>TF (AP2/ERF, C2C2, MYB, bHLH, NAC, HD-ZIP)<br>manganese transporter<br>phosphorylation (CAMK.SnRK3, S/T phosphatase.PPM/PP2C) | starch metab.<br>chll catab.<br>external stimuli (light, ETI network)<br>transition metal homeostasis<br>photophosphorylation (PSII, chlororespiration)<br>ABA (perception and sign.)<br>TF (AP2/ERF, MYB, bHLH, NF-Y)<br>phosphorylation ( S/T phosphatase.PPM/PP2C)<br>floral meristem identity control<br>phenolics (flavonoid biosynth.) | cell wall org. (pectin.rhamnogalacturonan I)<br>glycosyltransferase<br>ethylene<br>solute transp. (MFS superfamily.SP family (ERD6)) |
| MapMan | All | cell division (DNA replication)<br>call wall org. (cutin and suberin, cell wall prot.hydroxyproline glycoprot. activities)<br>ER fatty acid elongase. 3-ketoacyl-CoA synth.<br>glycosyltransferase<br>calcium dep. sign. (CBL-CIPK)<br>circadian clock (evening element reg)<br>ABA (conjugation, perception and sign.)<br>TF (AP2/ERF, MYB, bHLH, NAC)<br>phosphorylation (CAMK.SnRK3, S/T phosphatase.PPM/PP2C)<br>solute transp. (MFS superfamily (NRT1/PTR)) | cell wall org. (pectin, cell wall prot.hydroxyproline glycoprot. activities)<br>ER fatty acid elongase. 3-ketoacyl-CoA synth.<br>glycosyltransferase<br>circadian clock<br>external stimuli (light)<br>ABA (perception and sign.)<br>TF (AP2/ERF, MYB, bHLH)<br>phosphorylation (TKL kinase superfamily, S/T phosphatase.PPM/PP2C)<br>solute transp. (APC superfamily, MFS superfamily (NRT1/PTR)) | cell wall org. (pectin.rhamnogalacturonan I)<br>starch metab. (ADP-glucose pyroP complex)<br>glucosinolate biosynth<br>glycosyltransferase<br>phenolics (flavonoid biosynth., p-coumaroyl-CoA biosynth.)<br>solute transp. (MFS superfamily.SP family (ERD6))<br>TF (PLATZ) |
